# Neural dynamics for working memory and evidence integration during olfactory navigation in *Drosophila*

**DOI:** 10.1101/2024.10.05.616803

**Authors:** Nicholas D. Kathman, Aaron J. Lanz, Jacob D. Freed, Katherine I. Nagel

## Abstract

Working memory and evidence integration are fundamental components of cognition thought to arise from distributed circuits throughout the brain^1–2^. Theoretical^3,4^ and behavioral studies^5,6^ argue that both processes are required for plume navigation, an innate task in which animals use stochastic sensory cues to navigate towards the unknown location of an odor source^7–10^. Here we identify a small population of local neurons in the navigation center of *Drosophila*^11–13^ that exhibits both evidence integration and working memory dynamics during goal-directed olfactory navigation. Developing a closed-loop virtual plume navigation paradigm, we show that a bump of activity in this population ramps up with successive odor encounters, and can persist for variable intervals after odor loss. While bump activity persists, the fly maintains the goal heading it adopted during odor. Silencing these neurons impairs the persistence of upwind heading after odor loss. Simulations show that the time constant of persistence observed in these neurons optimizes navigation in a turbulent boundary layer plume. Our work localizes working memory and evidence integration to a specific group of genetically-identified neurons, which will facilitate the mechanistic dissection of these building blocks of cognition.

## Introduction

In many real-world navigation tasks, sensory information about the direction and location of a goal is only intermittently available. For example, an owl hunting a mouse by sound might get only sporadic and uncertain information about the location of its prey, or a leopard chasing an antelope by sight might have its view interrupted by brush. In such tasks, the ability to accumulate sensory information about the direction of a target over time, and to hold that information in working memory during loss of sensory input are critical to success. While the neural mechanisms underlying the processes of evidence integration and working memory have been studied extensively in non-human primates and rodents^1,2^, the distributed nature of these circuits has made it challenging to causally link neural activity patterns to behavior^14,15^.

Many animals use odor cues to navigate towards the unknown location of a food source^16,17^. Because odor plumes are turbulent, they generate stochastic cues that must be integrated and stored in memory in order to navigate effectively^17^. Behavioral studies in plumes argue that insects integrate odor encounters over time to drive navigational decisions^5,6^, while theoretical studies argue that an optimal searching agent should possess a working memory that allows it to continue moving in the goal direction during short blanks in the odor stimulus^3,4^. However, the potential neural substrate of these computations is unknown.

Recent evidence suggests that the fan-shaped body (FB), a part of the insect navigation center called the Central Complex (CX), is a key location for computing a goal direction during navigation. Output neurons of the FB can compare the fly’s current heading to a goal signal expressed in allocentric (world-centered) coordinates to steer the fly in a target direction^11,13,18^. Patterned activation of certain FB local neurons can drive goal-directed walking relative to a visual landmark^13^, or to the wind^19^. Wind direction is known to drive rotation of compass neurons (EPGs^20,21^), and can be flexibly tethered to visual landmarks through plasticity in the compass system^13,22^. These experiments suggest that the fly may use the wind direction as an allocentric reference frame during odor-guided navigation, and that FB local neurons can encode goal directions relative to this reference frame.

The FB is ideally positioned to play a role in odor-guided navigation. It receives odor information through a set of wide-field tangential neurons that are downstream of the associative learning center in the mushroom body^19,23–24^. Activation of some of these tangential inputs, and also of some FB local neurons, can drive orientation upwind^19,25^, while broad developmental disruption of the CX completely abolishes odor-gated wind orientation behavior^26^. However, the *in vivo* dynamics of FB goal signals and their precise role in natural navigation tasks are not well understood.

In a recent study, we identified FB local neurons labeled by the line VT062617-Gal4 whose sparse activation can drive walking in a goal direction relative to the wind^19^. Here we developed a closed-loop virtual paradigm to examine the activity of these neurons during odor-guided navigation. We find that these neurons exhibit a bump of activity that is activated by odor but persists after odor offset for variable intervals. While the bump remains active, we find that flies continue running in the direction adopted during odor, arguing that these neurons represent a short-term directional goal memory. By simulating a turbulent plume in closed-loop, we show that these neurons integrate olfactory evidence over a period of several seconds, a property not shared by a second parallel group of local neurons. Optogenetic silencing of these neurons impairs persistent directional running after odor loss. Simulating these dynamics in a model shows that a short-term memory for goal direction with a time constant similar to the one we measure in these neurons improves navigation in a turbulent boundary layer plume— a plume near a surface such as a walking fly would encounter. Our findings reveal internal dynamics that shape the transformation of stochastic sensory input into goal-directed navigation behavior.

## Results

### A persistent bump of activity correlates with maintaining a goal direction after odor loss

To investigate the activity of VT062617 neurons during on-going navigation, we developed a closed-loop virtual odor navigation paradigm. In this paradigm (Fig. 1a), the fly controls the direction of the wind in closed-loop, while a pulse of attractive odor (apple cider vinegar, 0.5% dilution in water) is delivered in open loop (see Methods). Similar to previous observations in both freely-walking^5,6^ and tethered flies^27,28^, we observed that flies walked more strongly upwind during odor compared to the pre-wind baseline period (Fig, 1b, Ext. Data Fig.1a). Odor-evoked trajectories were straighter than baseline walking (Ext. Data Fig. 1a), and biased towards, but not uniformly, upwind (Ext Data Fig. 1b). In some trials, flies showed a looping search-like behavior at odor offset, similar to search behavior evoked by odor loss in freely-walking flies^5,27,29^ (Fig. 1b, left-most trace). On many trials, however, flies maintained their trajectory for a variable number of seconds following the loss of odor (Fig 1b, right-most trace, Ext Data Fig. 1b,c). Straight trajectories after odor lasted significantly longer than those observed during the baseline period (Ext Data Fig. 1b,c).

**Figure 1:**
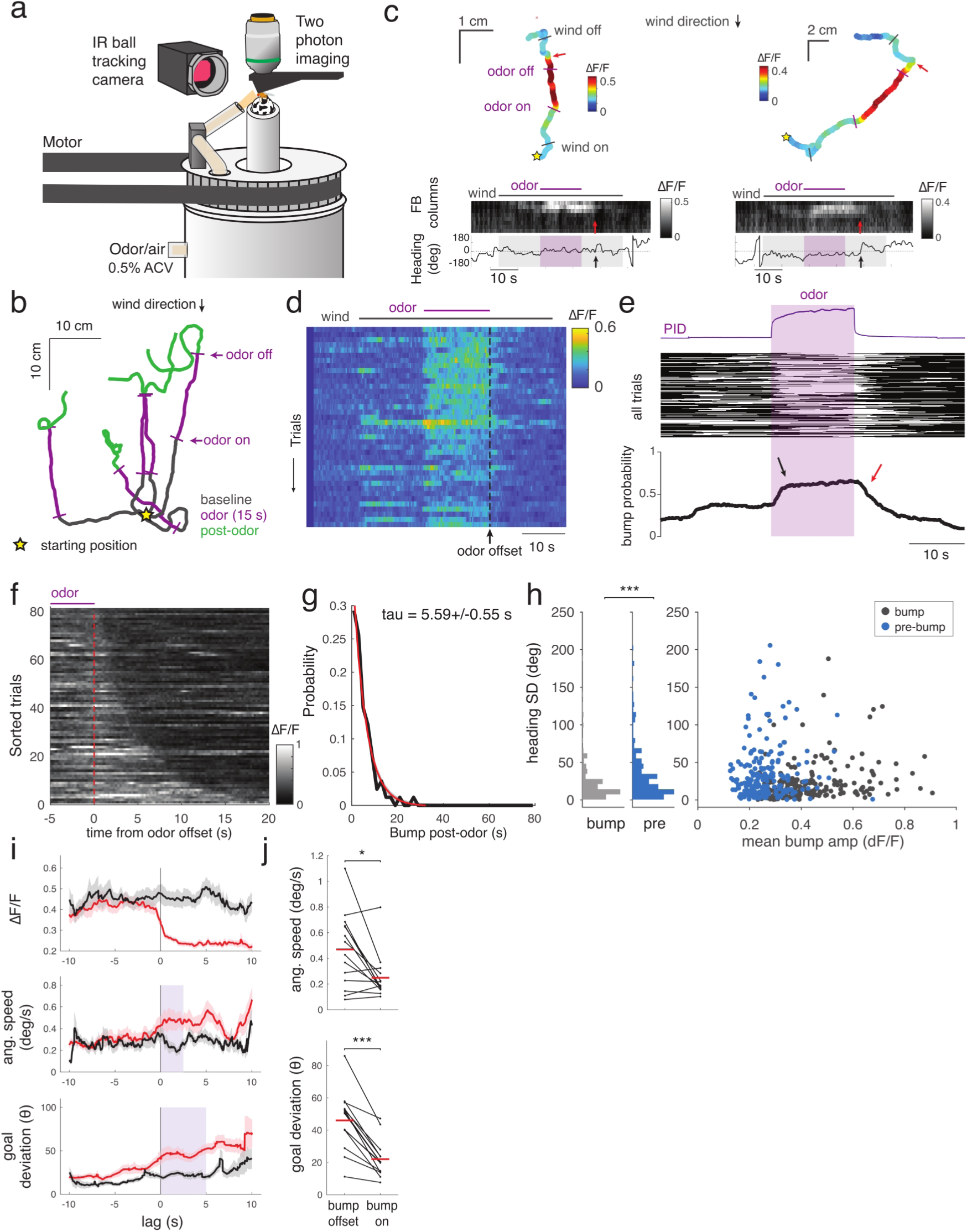
Persistent odor-evoked activity in VT062617 FB local neurons is associated with persistent goal-directed walking. **a**: Schematic of the preparation. A head-fixed fly walking on an air-supported ball can control wind direction (through a motorized rotary union) and odor (apple cider vinegar 0.5%) in open- or closed-loop. An IR camera tracks ball rotations. A 2-photon microscope images concurrent neural activity. **b**: Example trajectories from one fly performing closed-loop wind navigation. Odor evokes straight trajectories that are biased towards upwind directions. Yellow star: starting position, purple bars: odor ON, odor OFF. **c**: Examples of VT062617 activity during closed-loop wind with an odor pulse. Top: virtual trajectories, color-coded by bump amplitude (max ΔF/F across columns). Black bars indicate wind ON and wind OFF, purple bars indicate odor ON and odor OFF and red arrows indicates bump OFF. Note strongest activity during and after odor. Middle heatmap: activity across FB columns as a function of time. Wind and odor periods indicated above heatmaps, red arrow indicates time of bump OFF (left: 5.1s, right: 5.8s after odor OFF). Bottom: heading trace, 0° is upwind, black arrow indicates start of turn after straight walk **d**: Bump amplitude as a function of time for one fly across trials. Note robust responses to odor with variable offset timing, and more variable and transient responses to wind ON. **e**: Timing of bumps for all trials and flies (202 trials from 14 flies) relative to measured odor time course (PID— top). Note that bumps initiated during odor (black arrow) often persist after odor offset (red arrow). **f:** Normalized bump amplitude for all trials with bump activity that started during odor and ended after odor offset (n = 82 bumps from N = 14 flies), aligned at odor OFF and sorted by bump persistence duration. **g**: Probability distribution (black) of bump persistence durations shown in **f**. Exponential fit (dashed line) with confidence intervals (shaded in grey, tau = 5.59+/-0.55 s). **h**: Standard deviation of unwrapped heading (y-axis) versus bump amplitude (x-axis) for all bumps (222 bumps from 14 flies, blue) and for equivalent durations before each bump. Marginal distributions at left. Heading standard deviation during bumps is significantly lower than during pre-bump periods (paired t-test, p=5.0e-6). **i**: Trajectory features associated with bump offset in the post-odor period (red), compared to periods when the bump remained on (black). Lines show mean bump amplitude (top), angular speed (middle), and deviation from goal (bottom, see Methods) for n = 78 bumps from N=14 flies. Shaded regions represent S.E. across bumps. **j**: Both angular speed (top) and deviation from goal direction (bottom) increase significantly after bump OFF (comparison periods in shown as blue shaded regions in **i**, paired t-test, ang speed: p=0.02, goal dev.: p=0.0002).

Imaging from FB local neurons labeled by VT062617 (Ext Data Fig. 1d), revealed a localized bump of activity with dynamics related to both sensory input and ongoing navigation behavior (Fig. 1c). Bump activity was most strongly and consistently activated by odor (Fig. 1d,e, Ext. Data Fig.1e), weakly activated by wind onset in some trials (Fig. 1d,e), and was absent in interleaved trials without airflow (Ext. Data Fig. 1f).

Localized to a few columns (Ext. Data Fig.1g), bump activity was preferentially initiated during the odor period (Fig. 1e), but could persist for variable intervals into the post-odor period (Fig. 1d,e,f, Ext. Data Fig. 1h,i). The distribution of persistence times was well-fit by an exponential distribution with a time constant of 5.59 +/- 0.55 s (Fig. 1g). These data suggest that bumps are preferentially ignited by odor but can persist in the absence of odor through some kind of internal dynamics.

In several example trajectories, we observed that persistent bump activity in VT062617 neurons correlated with maintained straight movement (Fig. 1c). To test whether this observation held across trials and flies, we extracted all bumps from all trials and flies and plotted parameters of trajectories during bump periods compared to equivalent durations preceding each bump. We found that both the standard deviation of heading, and mean absolute angular velocity, were significantly reduced during bump versus pre-bump periods (Fig. 1h, Ext. Data Fig. 1j), consistent with the idea that bumps correlate with straighter movement. The presence of a bump correlated with reduced heading standard deviation during both odor and pre-odor periods (Ext Data Fig. 1k). Together, these analyses suggest that bumps turn on preferentially in response to odor and correlate with straight goal-directed movement as long as they remain active. If this is true, we would expect that persistent bump activity in the post-odor period should correlate with maintaining the goal trajectory adopted during odor. To test this hypothesis, we detected bumps that turned on during odor and persisted after odor offset (see Methods, Fig. 1i,j). We observed that while the bump remained on, trajectories remained close to the “goal trajectory” adopted during the odor period. In contrast, when the bump turned off, angular velocity transiently increased and trajectories deviated from this goal direction. We conclude that the presence of an activity bump in VT062617 correlates with maintaining a goal direction, including maintaining a goal for several seconds after the loss of odor.

If activity bumps in VT062617 neurons represent a goal direction, we would expect bump position to remain stable even when the fly turns^13,30^. Consistent with this hypothesis, we observed little movement of the bump during turns, and poor correlation with heading (Fig. 2a,b). We would further expect that this bump should remain stable when the wind is rotated, which should drive a change in the fly’s reference frame by rotating the representation in compass neurons^13,20,22^. When we rotated the wind during odor, we sometimes observed a compensatory change in heading, but this was not associated with any movement of the bump in VT062617 neurons (Fig. 2c,d). Compensatory turns could be driven by the difference between the rotated wind-anchored reference frame and the stable goal signal representing a direction relative to the wind^13,18^. We conclude that the activity bump in VT062617 most likely represents a short-term goal direction relative to wind that can be evoked by odor and persists after odor loss.

**Figure 2:**
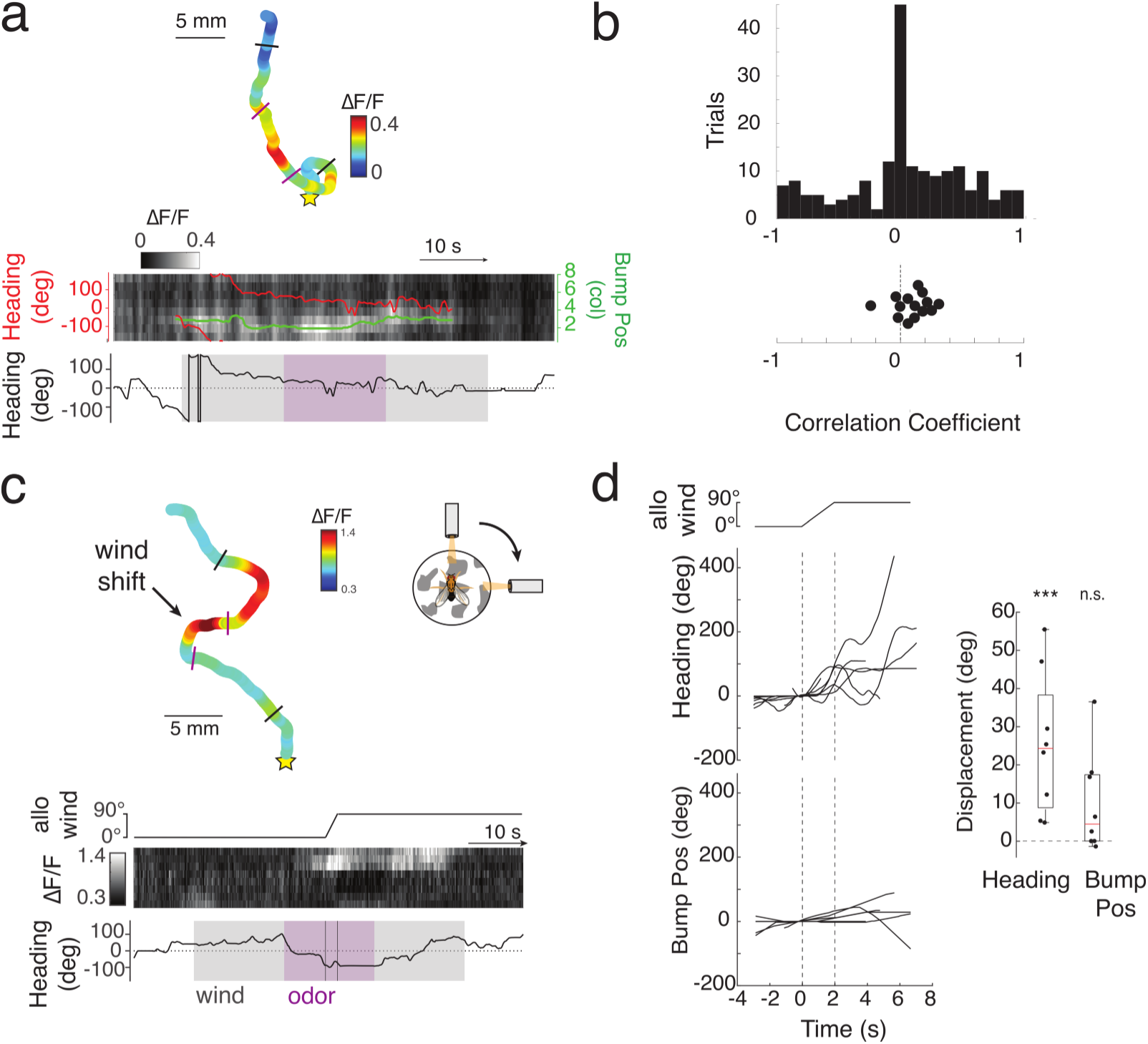
Bump activity in VT062617 neurons is stable during changes in heading. **a**: Example trial where bump position does not change with changes in heading. Top: fictive 2-D trajectory color-coded by bump amplitude; symbols as in Fig. 1c. Bottom: Heatmap shows activity across columns with bump position (green) and heading (red) overlaid. Heading for whole trial shown below. **b**: Distribution of correlation coefficients between bump position and heading for all trials from all flies (top, n=196 trials) and averaged for each fly (N=14 flies). Pooled data were not significantly different from zero (one-sample t-test, mean=0.043, p=0.220). Fly averages showed a very small positive correlation (mean=0.095, p=0.03). **c**: Response of a single fly to a 90° wind shift during odor. Top: fictive 2-D trajectory color-coded by bump amplitude; symbols as in Fig. 1c. Bottom: allocentric wind direction (top), VT062617 activity across columns (middle), and heading (bottom). Note stability of bump position during heading deviation following wind shift. **d**: Left: Heading response to open-loop wind shift in 8 trials from 4 flies that responded significantly to the wind shift (see Methods). Dashed lines represent start and stop of shift. Middle: Bump position for the same trials shown to the left, offset to an initial position of 0. Note lack of consistent movement. Right: Displacement of heading and bump position for the 8 trials shown. Heading displacement is significantly different from zero (mean: 25.41°, one-way t-test: p=0.006) while bump displacement is not (mean: 9.87°; one-way t-test: p=0.072).

### Integration of olfactory evidence during turbulent plume navigation

In natural environments, odors form turbulent plumes in which odor fluctuates stochastically as a function of distance and direction to the source^31–34^. Therefore, we next sought to examine the activity of VT062617 neurons as flies navigated a turbulent plume. To implement a virtual plume environment, we leveraged a previously recorded movie of odor concentrations in a boundary layer plume at approximately fly height and windspeed comparable to our experiment^34^ (Methods, Fig. 3a). As the fly navigated in 2D, we used time and the virtual position of the fly to index into this movie and determine the odor concentration at the virtual location of the fly. We then used an adaptive compression algorithm^5^ that mimics aspects of peripheral olfactory encoding^35^ to binarize this concentration relative to an adaptive threshold, and delivered the resulting odor pulses to the fly. As a result of this algorithm, a fly navigating our virtual odor plume encounters a stochastic series of odor pulses that increase in frequency as the fly approaches the virtual source.

**Figure 3:**
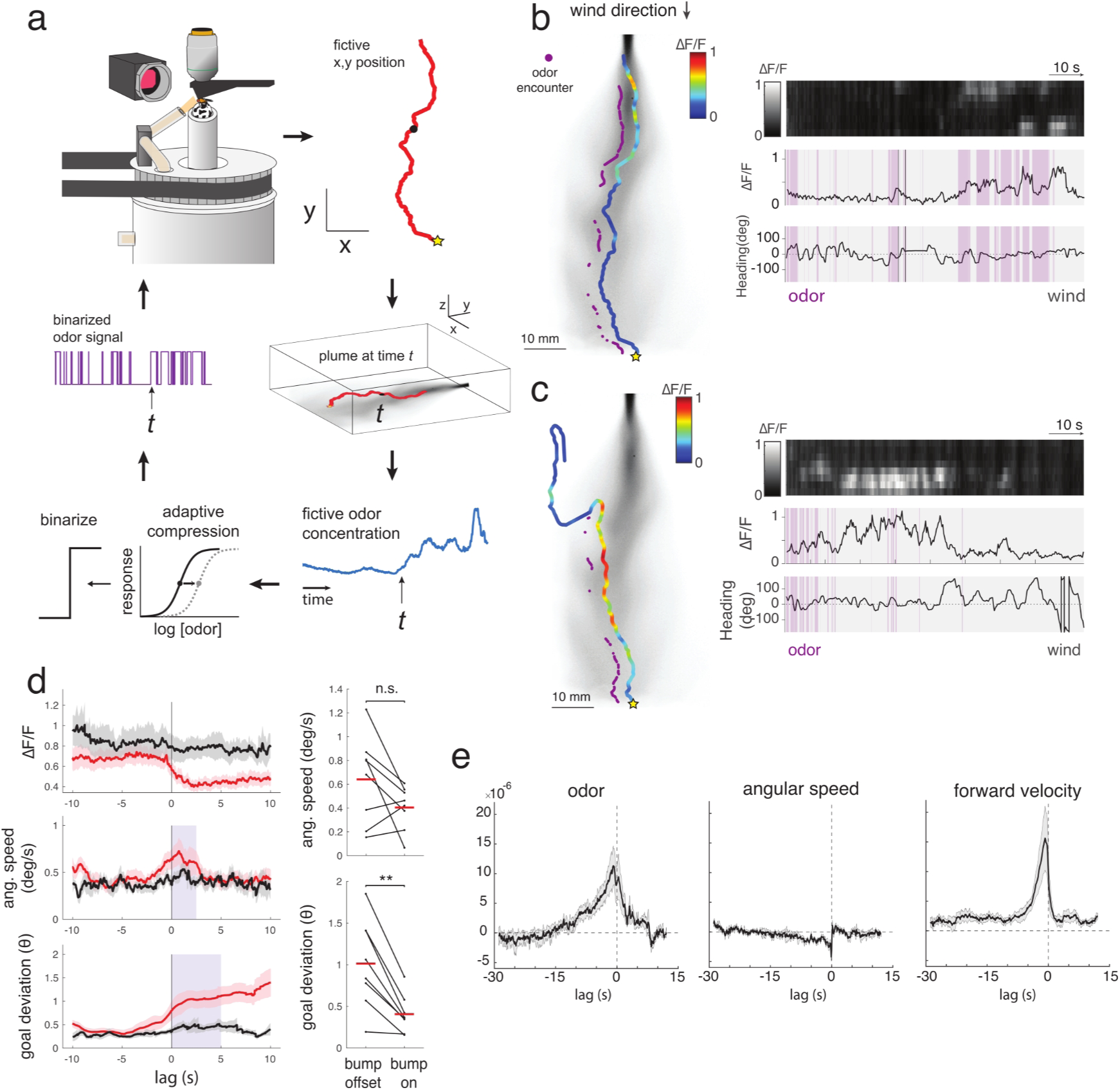
Evidence integration dynamics during closed-loop plume navigation. **a**: Closed-loop plume navigation paradigm. Time and fictive fly position are used to index into a previously recorded movie of plume dynamics. Measured odor concentrations are converted into binary odor signals based on an adaptive compression algorithm (see Methods). The frequency of the resulting odor pulses depends on location and distance from the source. **b**: Example trial from a fly navigating the virtual plume. Left: virtual 2D trajectory color-coded by bump amplitude, superimposed on a still of the plume movie. Purple dots indicate fictive locations at which odor was presented. Right: activity across FB columns (top), bump amplitude (middle), and heading as a function of time (bottom). Purple bars indicate odor periods. Note minimal responses to early odor presentations and gradual ramping of activity over time. **c**: Trajectory and activity for another example trial. Note prolonged period of activity associated with straight walking. **d**: Left: Analysis of locomotor features around the bump OFF period (red) compared to the bump ON period (of equal length, black) during plume navigation Left: Analysis of locomotor features around the bump OFF period (red) compared to the bump ON period (of equal length, black) during plume navigation (n = 143 bumps from N= 8 flies). Shaded regions represent S.E. across bumps. Right: Goal deviation was significantly higher over the time window shaded in blue (left) while angular speed was not (paired two-sample t-test, ang. speed, p = 0.12, goal dev: p = 0.01, N=8 flies). **e**: Linear filters relating neural activity to odor, angular speed, and forward velocity, each mean +/- S.E. for N=6 flies. Odor and forward velocities have an integrating shape with a longer integration time for odor than for forward velocity. Angular speed filter is negative indicating a correlation with decreased angular speed (straight walking).

Plume navigation revealed additional dynamics in VT062617 neurons. As the fly navigated the plume, we observed initially weak responses to odor that grew with repeated odor encounters (Fig. 3b). We also observed instances where a series of odor encounters ignited a long-lasting bump of activity, that was again associated with maintaining a straight trajectory (Fig. 3c). To ask whether bump activity correlates with a maintained trajectory in this more complex environment, we computed parameters of fly trajectories associated with maintained bump activity after odor offset (Fig. 3d). Similar to our observations with isolated odor pulses, we found that persistent bump activity was associated with lower goal deviation, and that bump offset was associated with a significant increase in deviation from the goal direction adopted during odor.

During our closed-loop navigation tasks, odor, locomotor parameters, and navigation behavior are all correlated with one another, making it challenging to disentangle the contribution of each parameter to the signal observed in VT062617 neurons. To address this issue, we took advantage of the stochastic nature of odor encounters in our plume paradigm to compute decorrelated linear filters between bump amplitude and odor, forward velocity, and angular speed. Adopting techniques from the sensory coding literature^36^, we computed the correlation matrix between these variables and applied a regularized pseudo-inverse of this matrix to the cross-correlogram of each parameter and bump amplitude (see Methods) to compute decorrelated filters relating odor and behavior to neural activity. This analysis allows us to examine the independent contribution of odor and behavior to bump activity, despite the fact that these variables are correlated in our dataset. Our decorrelated filter analysis reveals that both odor and behavior contribute to bump activity, with distinct dynamics. The filter describing the relationship between odor and neural activity (Fig. 3e, Ext. Data Fig. 2a) had an “integrating” shape with a single peak that extends ∼10 seconds into the past, supporting the idea that these neurons integrate odor information over several seconds. In contrast, the filter relating angular speed (turning) to activity was entirely negative (Fig. 3e, Ext. Data Fig. 2b), consistent with the observation that activity is strongest during straight walking. We also observed a positive integrating filter for forward velocity (Fig. 3e, Ext. Data Fig. 2c), although this filter was narrower than the filter for odor. This forward velocity filter supports observations during odor pulse trials, where bump onset was associated with an increase in forward velocity, and forward velocity was higher when a bump was present (Ext. Data Fig. 2d,e). Our plume experiments thus reveal slow temporal integration of odor input, and support our previous finding that persistent activity in VT062617 neurons is associated with faster, straighter trajectories.

### Persistent activity and odor integration are not widespread in the FB

The dynamics we observe in VT062617 neurons might arise within these neurons or might be more broadly shared across the fan-shaped body. To address the specificity of these dynamics, we imaged from a second line, 52G12-Gal4, that labels a large set of ventral FB local neurons (Ext. Data Fig. 3a,b). Imaging from 52G12 neurons in the plume paradigm revealed transient responses to repeated odor encounters without the gradual ramping and persistence we observed in VT062617 neurons (Fig. 4a). The filter relating odor to activity for 52G12 was much narrower than the filter for VT062617, indicating a shorter integration time (Fig. 4b). The filter relating angular speed to activity had a positive peak for 52G12, in contrast to the negative peak for VT062617, indicating a correlation with turning rather than straight walking (Fig. 4c).

**Figure 4:**
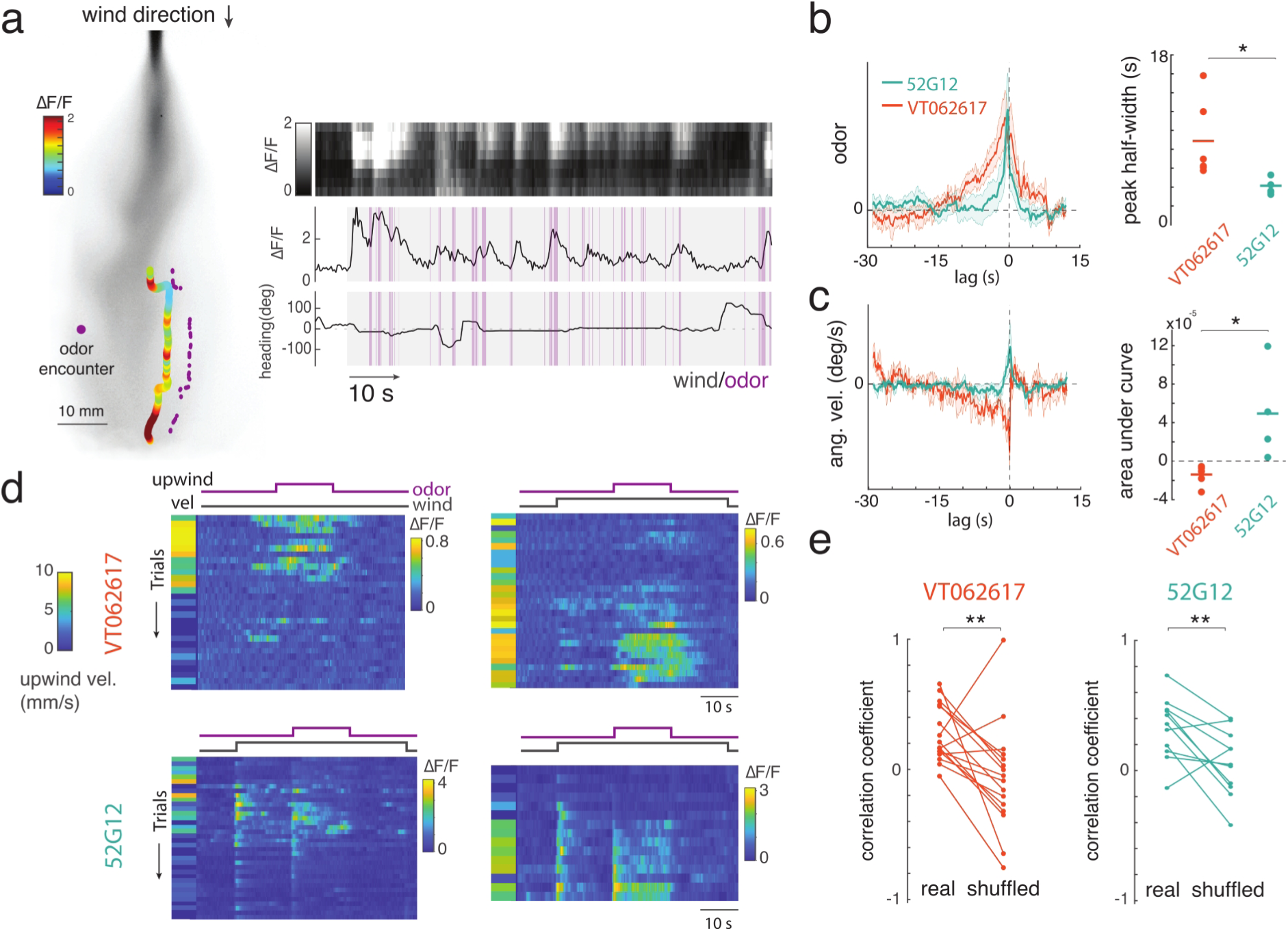
Distinct local dynamics and shared slow dynamics in a population of ventral local neurons. **a**: Example trial from a fly expressing GCaMP in 52G12 neurons navigating the virtual plume. Left: trajectory color-coded by bump amplitude. Right: activity across FB columns (top), bump amplitude (middle), and heading (bottom) as a function of time. Purple bars indicate odor periods. **b**: Linear filters relating odor to activity in 52G12 neurons (green, mean +/- S.E. from N=5 flies) and VT062617 neurons (orange, replotted from Fig. 2). 52G12 filters are significantly narrower than VT062617 filters (two-sample t-test, p=0.03). **c**: Linear filters relating angular speed to activity in 52G12 versus VT062617 neurons. 52G12 filters have a positive area under the curve, indicating a correlation with turning, while VT062617 filters have a negative area (two-sample t-test, p=0.015). **d**: Example data from 4 flies (top: VT062617, bottom: 52G12) showing upwind velocity during odor (color bar, left) and bump amplitude (right) across trials. **e**: Trial-by-trial correlation coefficients between upwind velocity and total bump amplitude for each fly are significantly higher than for trial-shuffled control data (paired t-test, p-values: VT062617, p=0.001; 52G12, p=0.013).

Consistent with these observations, we found that 52G12 responses to single odor pulses were transient rather than persistent (Ext. Data Fig. 3c,d), and correlated with turning rather than straight walking (Ext. Data Fig. 4e). Thus, the coding features of VT062617 neurons— gradual integration of olfactory evidence and persistent activity associated with straight trajectories— are not shared across the FB, but are specific to a subset of local neurons.

While neurons labeled by 52G12 and VT062617 showed distinct local encoding dynamics, both showed similar modulation on the slower timescale of trials (Fig. 4d). Many flies walked upwind only on a subset of trials in an experiment. Runs of trials with high upwind displacement could occur both early and late in the experiment for both genotypes, arguing that lack of engagement does not reflect simple sensory habituation. We observed greater activity in both VT062617 and 52G12 neurons during periods when the fly showed strong upwind displacement. Upwind displacement and total bump activity were positively correlated across both genotypes (Fig. 4e), significantly greater than in shuffled trials. These data indicate that while different FB local neurons track distinct local features of odor dynamics and self-motion, activity across the FB may be co-regulated by behavioral state on slower timescales.

### VT062617 neurons play a causal role in persistent goal-directed navigation

Our plume experiments argue that VT062617 neurons integrate odor information over time and maintain this information for seconds after odor offset. We therefore wondered if these neurons contribute causally to persistent goal-directed navigation behavior. We first asked whether freely-walking flies exhibit similar persistent trajectories to those we observed in our virtual paradigm. We presented freely walking flies in a laminar wind tunnel with variable numbers of odor pulses at 1Hz, and examined the dynamics of their orientation behavior (Fig. 5a-c). We found that increasing numbers of odor pulses led to a greater probability of upwind walking, that saturated around 4 pulses (Fig. 5b,c). Upwind walking persisted for 5-10 seconds after the end of the last odor pulse, and decayed with a uniform time course across pulse numbers (Fig. 5b). These results indicate that persistent upwind walking that outlasts the odor stimulus also occurs in freely-walking flies.

**Figure 5.**
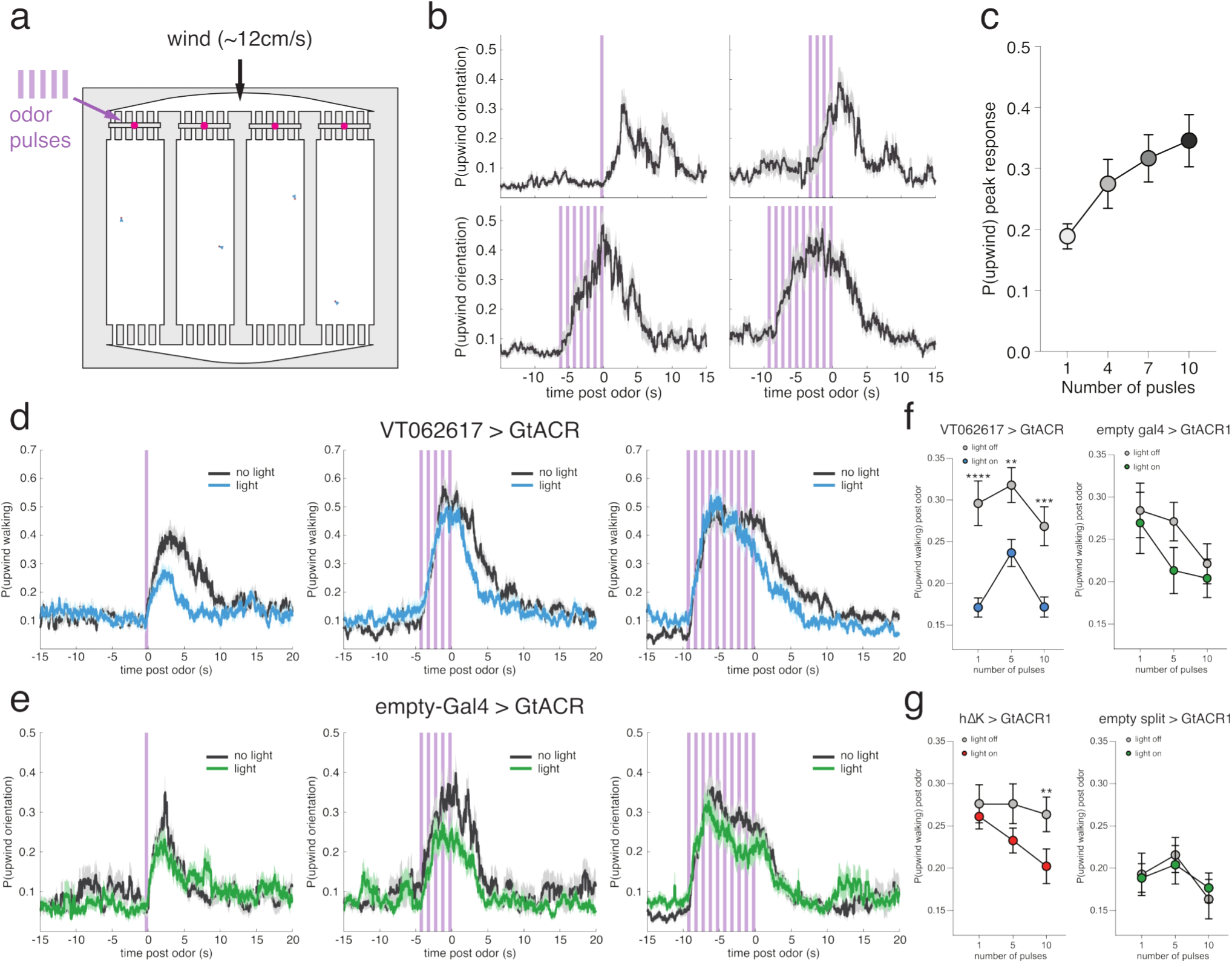
VT062617 neurons are causally required for persistent upwind navigation. **a**: Schematic of behavioral apparatus used to track freely moving flies, showing odor and wind inputs. **b**: Probability of upwind walking as a function of time, for different numbers of odor pulses (500 ms each) in wild type flies. **c**: Mean probability of upwind walking during peak response period (2-6 s, relative to odor start) for each pulse number. Data are mean +/- S.E. across flies. **d**: Probability of upwind walking in VT062617>GtACR flies responding to 1, 5, or 10 pulses of odor with light OFF or light ON. Note decrease in upwind walking probability after odor OFF on light ON trials across stimuli. **e**: Probability of upwind walking in empty-gal4>GtACR flies responding to the same stimuli with light OFF and light ON. **f**: Left: Mean probability of upwind walking during the 10 s post-odor period in VT062617 > GtACR1 flies during light ON (blue) is significantly lower than in light OFF (grey) trials. Right: same comparison, but with empty-gal4 > GtACR1 flies during light ON (green) and light OFF (grey) trials. **g**: Left: Mean probability of upwind walking during the 10 s post-odor period in hΔK split > GtACR1 flies during light ON (red) is significantly lower than in light OFF (grey) trials after 10 pulses. Right: same comparison, but with empty-gal4 > GtACR1 flies at the same light level during light ON (green) and light OFF (grey) trials. All statistics shown in Ext Data Table 5.

To ask whether VT062617 neurons play a role in these dynamics, we presented flies expressing the optogenetic silencing agent UAS-GtACR1^37^ in VT062617 neurons with variable numbers of odor pulses. We measured upwind walking in interleaved trials with blue light that silences the neurons, and in the absence of light. We found that silencing VT062617 neurons led to reduced persistence in upwind walking across pulse numbers (Fig. 5d). The probability of walking upwind in the 10s period after odor OFF was significantly reduced for all stimulus configurations (Fig. 5f). In genetic control flies, light had no effect on the persistence of upwind walking (Fig. 5e,f). To ask if this effect is specific to fan-shaped body neurons labeled in this line, we repeated our experiment using a split-Gal4 line specific for hΔK neurons^38^, the dominant FB cell type in VT062617^26^, to drive GtACR1. We also observed a decrease in persistent upwind running using this split-Gal4 line, although a higher light level was required to obtain a behavioral effect and we only observed a significant reduction with the longest pulse train (Fig. 5g, Ext Data Fig. 4a,b). We conclude that FB local neurons labeled by VT062617 neurons contribute causally to maintaining a trajectory after odor loss, but that other neurons are likely to contribute to the initial selection of a goal trajectory. We note that wind speeds in our freely walking wind tunnels (∼12 cm/s) are a subset of those we used on the walking ball, and that higher wind speeds promoted more effective upwind tracking (Ext. Data Fig. 4c). Wind speed was consistent throughout wind tunnel trials^5^ and during odor pulses on the walking ball (Ext Data Fig. 4d).

### A persistent goal representation improves navigation in turbulent environments

Our data argue that VT062617 neurons are not required for a fly to select an initial upwind trajectory in response to odor, but that their activity allows the fly to maintain its trajectory for some time after odor loss. Intriguingly, this type of working memory was recently predicted by two theoretical studies of plume navigation using purely computational methods^3,4^. Both studies found that an optimal agent should integrate odor information over a window related to the average duration of blanks in an odor plume, allowing the agent to determine whether a gap in odor is expected (meaning the agent is within the plume envelope) or unexpected (because the agent has lost the plume). To test whether the dynamics we observed in VT062617 neurons could perform such a function, we integrated previous models of fly olfactory navigation^5,29^ with the finite state controller framework of Verano et al.^3^ In this model, the fly can adopt one of three navigational states: baseline walking, goal-directed upwind walking, or search. Critically, the transition from goal-directed behavior to search is probabilistic with a time constant *τ _p_*, allowing the system to remain in the goal-directed state for variable intervals after odor offset (Fig. 6a,b, Methods). We fit coefficients of the model to approximately reproduce the distribution of forward and angular velocities measured in our walking ball paradigm (Fig. 6c). In particular, the mean forward and angular velocity of the model and real data overlapped to within 95% confidence intervals (shaded regions in Fig. 6c).

**Figure 6.**
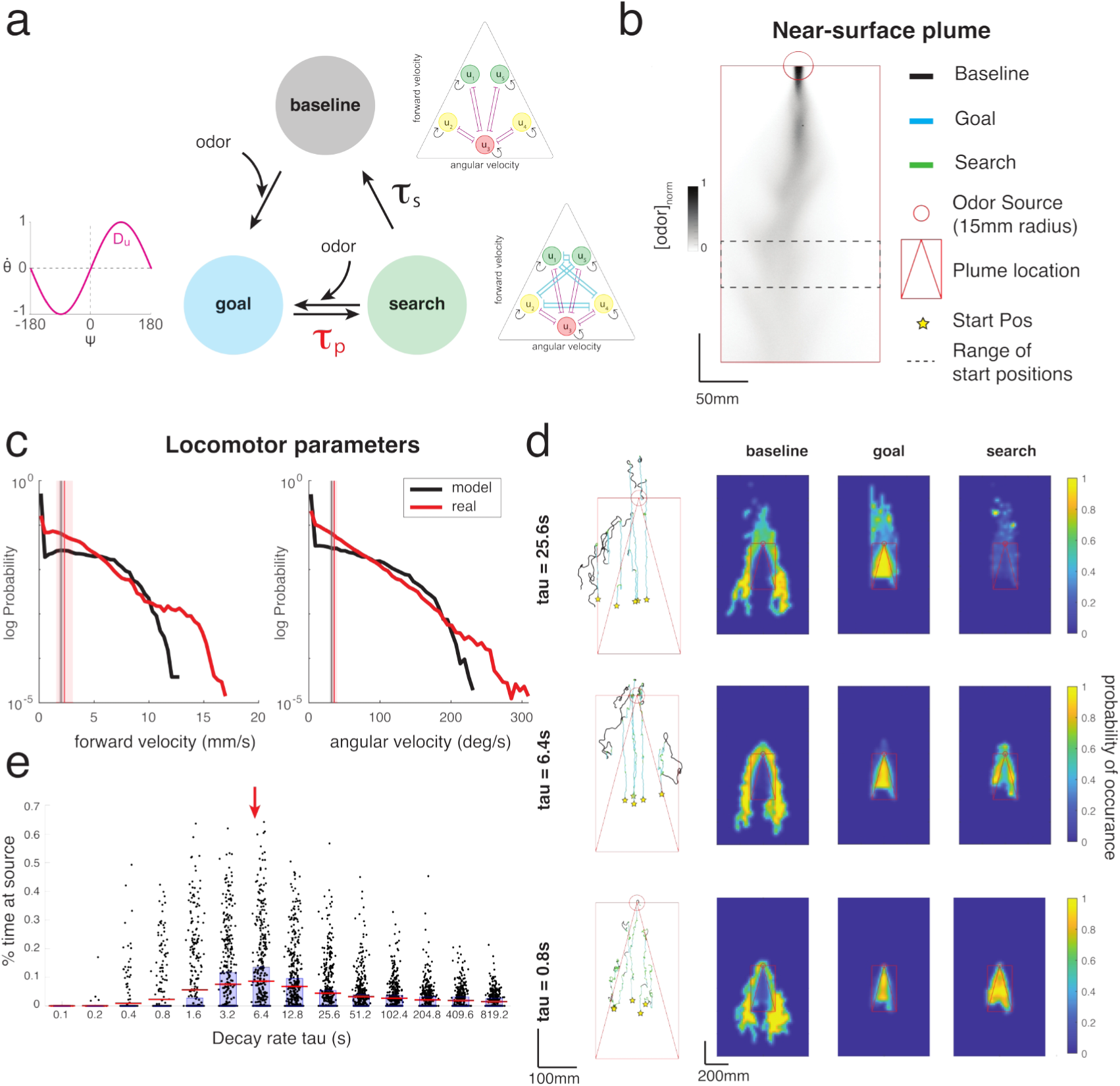
Measured persistence times closely match optimal memory durations for a walking fly navigating a boundary-layer plume. **a**. Schematic of the state transition model. The agent switches between three behavioral states, baseline (gray), goal-directed upwind walking (blue), and local search (green) based on odor inputs and internal probabilistic dynamics. Transitions between the goal state and the search state are governed by a tunable time constant *τ _p_*. **b.** Still frame of a boundary layer plume as the simulation environment. Dashed black rectangle is starting zone for each trial. Symbols define graphical elements used in subsequent panels: yellow start is trial start location, red square and triangle is the plume sampled area and odor envelope estimate, red circle is the “success” area for finding the source location; colored lines are state annotations for example trajectories. **c**. Distributions of locomotor parameters (forward and angular velocity) for real flies in our walking ball data (red) and simulated agents (black) during baseline walking. Real (red) and model (gray) means shown as vertical lines with +/- 95% confidence interval shaded. Forward vel, real: 2.3+/-0.77mm/s, model: 1.99+/-0.2mm/s. Ang vel, real: 35.5+/-4.25 deg/s, model: 31.7+/-2.28 deg/s. **d.** Example trajectories from the model (left) and state-specific occupancy maps (right) showing positions during baseline (top), goal (middle), and search (bottom) states for three different values of *τ _p_* (rows). At the optimal *τ _p_*, the agent adopts that goal state at most locations within the plume envelope and the search state is concentrated at the edges of the plume envelope. **e.** Success rate (see Methods) as a function of *τ _p_* for 500 runs of the model. Peak performance occurs at an intermediate *τ _p_* value 6.4s. Red line is the mean, blue box shows middle quartiles. Red arrow shows measured τ_real_ = 5.59 s from VT062617 imaging.

We ran this model in the same previously recorded odor plume we used for our plume experiments and varied the time constant *τ _p_* of memory for the goal state (Fig. 6d,e). When this time constant was small (Fig 6d, bottom row), we found that flies began searching during brief pauses in odor within the plume.

Consequently, they often remained trapped near the base of the plume and did not make strong progress upwind. As the time constant increased (Fig 6d, middle row), flies made more effective upwind progress, and search behavior became concentrated at the edges of the plume envelope. At longer time constants, flies ran past the source, and search success decreased (Fig 6d, top row). Although it is not known what causes flies to stop at a food source, our model suggests that very long integration times could be deleterious when searching a stochastic signal. Thus, consistent with previous studies, we find that an optimal memory time constant allows flies to integrate stochastic odor encounters within the plume, and correctly localize search behavior at the edges of the plume envelope. Strikingly, the optimal memory time constant for our model was very similar to time constant of persistence we observed in VT062617 neurons (Fig. 6e, red arrow). We conclude that the dynamics of persistent activity observed in these neurons provide an optimal integration of olfactory information for a fly navigating a turbulent plume.

## Discussion

### A persistent goal representation supports navigation with sparse sensory input

During natural navigation tasks, sensory cues indicating the direction or location of a target can be sparse. Turbulent plumes epitomize this type of task, as odor encounters in a plume are intermittent, with statistics that depend in complex ways on medium, windspeed, lateral and longitudinal distance from the source, and distance from the ground, among other factors^31–34^. Ideally, an agent trying to find the source of a plume should maintain its goal heading if it is within the plume envelope, and initiate search or recovery behaviors only when this envelope has been lost^3,4,17^. However, instantaneous measurements of odor concentration are insufficient to discriminate between expected gaps due to turbulence and gaps due to plume loss^37^.

One solution for an animal navigating a stochastic environment is to integrate sensory information over time. By integrating odor input as it moves, a fly could generate an internal estimate of whether it is in the plume or not, and use this to more effectively determine whether it should continue pursuing its goal, or initial search behaviors^5,6,17^. However, to be effective, such a signal should be tuned to the expected duration of gaps within a plume.

Here we identify a neural goal signal that exhibits integration and persistence tuned to the properties of odor plumes that a walking fly is likely to encounter. In a previous study, we showed that sparse activation of this population of neurons can drive movement in a reproducible direction relative to the wind, suggesting that it can encode a navigational goal^19^. Here we show that this population exhibits a localized bump of activity that turns on in response to odor and remains on while the fly maintains a goal heading relative to wind. Rotation of the wind does not cause a rotation of the bump, consistent with the idea that it represents a goal direction relative to a reference frame anchored by wind, rather than a heading estimate^13,30^. Silencing of these neurons curtailed persistent directional running after odor offset, arguing that they play a causal role in maintaining a goal direction. Interestingly, the persistence time of these neurons appears to be probabilistic, rather than fixed (Fig. 1d,f). Simulations with a similar probabilistic persistence time showed effective integration in a boundary layer plume, arguing that the dynamics we observed *in vivo* are well-suited to navigating the types of environment a walking fly is likely to encounter. In our imaging data, different flies showed different mean persistence durations (Ext Data Fig. 1i), suggesting that this integration time might be plastic. Future experiments might explore whether the duration of persistence in these neurons can change depending on the statistics of the odor environment or the fly’s mode of locomotion.

Recent studies of the FB suggest that it translates variables into allocentric coordinates^13,18,39,40^, and can represent goal directions for navigation in this world-centric system^13,19,30^. An allocentric representation of estimated source direction could allow the fly to weigh and accumulate evidence for or against a particular direction in world coordinates, rather than having to update these representations with each turn. Behavioral experiments argue that insects integrate many different cues to navigate towards an odor source, including the temporal pattern of odor encounters^41^, differences in odor concentration across the antennae^42,43^, the motion of the odor itself^44^, and the direction of the wind^45^. A centralized goal signal could allow these diverse cues to be integrated into a single estimate of goal direction. The direction signal we identified could also be inverted, perhaps through the system of hΔ neurons that translates signals by 180° across the array of the FB, in order to allow a fly to relocate the edge of an odor after leaving it^46,47^. Future experiments could investigate the integration of convergent and divergent cues to source location at the level of the signal we describe.

### Working memory and evidence integration in a genetically tractable central brain circuit

The persistent goal signal we characterize here shares properties with working memory signals described in vertebrate systems^2^. Classical experiments identified persistent stimulus-tuned activity in the prefrontal cortex of monkeys as a potential neural correlate of working memory^48–49^, while ramping activity in several cortical and subcortical regions has been linked to integration of noisy sensory evidence^1,14,50^.

Dominant models of working memory in vertebrates rely on attractors generated by recurrent excitatory connectivity^2,15,51,52^, but direct evidence for these networks has been difficult to obtain due to the distributed nature of vertebrate circuits^14,15,51,52^. The major local neuron type labeled by VT062617 is hΔK^26^, which shows strong recurrent connections with another CX cell type known as PFG^12^. In a follow-up paper, we investigate the dynamics of these two populations using specific split-Gal4 lines, and show that recurrent connections between them are critical for the formation of persistent bump activity^53^. The identification of working memory and evidence accumulation dynamics in genetically identified neurons with complete connectome information^12^ opens these processes to mechanistic dissection at a cellular and synaptic level. A major question going forward is to what extent dynamics can be predicted from patterns of recurrent connectivity. For example, do the more transient dynamic observed in 52G12 reflect weaker recurrent connectivity in these neurons?

Understanding how different species represent goals for navigation remains an area of active research^54–56^. *Drosophila* larvae, which navigate in much more stable odor gradients^57^, lack a fully developed CX^58^, and may require only egocentric computations to navigate. While allocentric representations of goal direction are now established in adult flies^13^, it remains unclear if the FB also encodes goal locations. A recent theoretical model has suggested that insects could use a pair of FB vectors, one pointed towards home, and one estimating a goal direction, to triangulate the location of a goal^59^. Recent evidence suggests that flies can learn the location of a goal relative to olfactory landmarks^60^. To determine whether the signal we have identified in VT062617 represents a goal location or a goal direction will require experiments in which the fly receives specific cues to goal location. In contrast to flies, rodents foraging for odorous food rapidly switch from using odor cues to remembered spatial positions^61^. In rodents performing odor-guided localization tasks, mixed odor and spatial representations develop at many places along the pathway from odor input to hippocampus^62,63^. How the distinct navigation systems of vertebrates and invertebrates might enable different behaviors remains an open question^64^. Using ethological tasks to tease apart the nature and causal role of goal representations can provide insight into the structure and organization of navigation circuits across species.

## Methods

### Animals

#### Fly strains and husbandry

All flies were raised at room temp (approximately 24°-26°C) on a 12 hour light/dark cycle on standard cornmeal-agar food. For imaging experiments we used 7-12 day old female flies that were starved for ∼24 hours. Experiments were run from 0-9 hours ZT. For freely walking wind tunnel experiments, we used 3-7 day old female flies. Flies were light-shifted for at least three days and then starved for 18-24 hours before experimentation. For GtACR1 flies, fly food was supplemented with hydrated potato flakes containing all-trans-retinal (35mM stock: Sigma, R2500, dissolved in ethanol, stored at −20°C).

Genotypes used in each figure were as follows:

|  |  |
| --- | --- |
| Figure 1-4, Ext. Data 1,2 | VT062617-Gal4 > UAS-GCaMP7f,UAS-tdTomato |
| Figure 3, Ext. Data 3 | 52G12-Gal4 > UAS-GCaMP7f,UAS-tdTomato<br>VT062617-Gal4 > UAS-GCaMP7f,UAS-tdTomato |
| Figure 5, Ext. Data 4 | w<sup>1118</sup> 5905 (5a-c)<br>VT062617-Gal4 > UAS-GtACR1 (5d,f)<br>Empty-Gal4 > UAS-GtACR1 (5e,f)<br>VT062617AD;VT037223DB > UAS-GtACR1 (5g, Ext. Data 4b)<br>EmptySplit-Gal4 > UAS-GtACR1 (5g, Ext. Data 4b)<br>VT062617-Gal4 > UAS-GCaMP7f,UAS-tdTomato (Ext. Data 4c,d) |

### Experimental Procedures

#### Closed-loop olfactory navigation paradigm

We designed a wind and odor delivery system in which the angular position of a wind tube and the status of a binary odor valve could be controlled in either open- or closed-loop by parameters of the fly’s locomotion, as measured on an air-supported walking ball. Wind and odor delivery was accomplished via a rotary union (Dynamic Sealing Technologies, # LT-2141-OF-ES12-F1-F2-C2) with three isolated channels that allowed continuous rotation of wind and odor outputs while maintaining isolation and pressure. The three channels carried wind, odor, and compensatory air for the odor OFF condition so that equal airspeed was maintained with and without odor. Air to all lines first passed through a pressure regulator (Cole-Parmer MIR2NA), and charcoal filter (Drierite 26800 Drying column, with desiccant replaced by activated charcoal). The wind line was then humidified and passed through a mass flow controller (Aalborg Model GFC17), and set to one of three flow rates: 0.2, 0.4, or 0.6 L/min. Air to the odor (0.5% apple cider vinegar) and comp (humidified air) lines was set to a flow rate of 0.3 L/min using a flowmeter (Cole-Parmer PMK1-010608). Air from the wind and odor/comp lines were mixed in a customized 3D-printed chamber that led to a 3.5cm Teflon laminarizing tube (5mm ID) extending to 2 cm from the fly’s antennae. Wind, comp, and odor entered the mixing chamber sequentially to prevent back flow of odor into clean air lines.

The wind ON condition was achieved by opening a solenoid valve (Cole-Parmer Masterflex P/N 01540-11) on the wind line and a high-speed three-way solenoid valve (Lee company, LHDA1233115HA) to divert air through the comp line rather than the odor line. In the ODOR ON condition air was instead diverted through the odor solution. All three valves were positioned immediately before the rotary union. Wind and odor lines were of Tygon tubing of either 1/16” ID or 1/8” ID.

Fly behavior was monitored using a lightweight walking ball (9 mm diameter General plastics FR-7110), supported by air from a custom 3D-printed holder. Airflow supporting the ball was controlled by a flow meter (Cole-Parmer PMK1-010608) at ∼0.4 L/sec. The ball was painted with spots of black ink for tracking by the FicTrac sphere tracking software^1^ using a Point Grey Camera (100 fps, Grasshopper, 94mm/0.5x InfiniStix Proximity Series lens, Edmund Optics Focal Length Extenders, working distance=15cm) and illuminated using two IR LED fiber optic cables (Thorlabs M118L03 flat cleave patch cables, M850F2 fiber coupled LEDs, and LEDD1B drivers). Two secondary cameras (30fps, FLIR Blackfly, Computar 8.5mm 1:1.3 adjustable lens) viewed the front and side of the fly for positioning.

The fly was held in place using a custom 3D-printed holder made of opaque resin (Grey Resin V5, Formlabs), with a tapered saline well that opens to a .9 x .9 mm imaging window at its nadir. Resin thickness around this window is necessarily thin (∼25-50 microns) to accommodate the objective’s working distance to the exposed brain below the window. The holder was held in place onto a fixed steel platform using a magnetic mount (Siskiyou MGB/20).

#### Closed-loop wind position from a walking ball

During closed-loop wind delivery, we used a microcontroller (TeensyDuino +USB) and stepper motor (Oriental PKP566FMN24A, with controller CVD524-K) to rotate the rotary union on which the wind tube was mounted, as shown in Figure 1A. The amount of rotation per 10ms interval was set to be equal to the rotation around the z axis of the walking ball. Wind tube position was updated based on a running average of measured z-rotations over 10 samples at 100 samples/s. Z-rotations above 1000°/s were clipped. In addition, the motor controller filtered out some rapid changes, leading to small amounts of error accumulation (∼0-90° per trial, depending on the amount of rapid turning). Therefore, an IR beam-break (Adafruit #2168) was used to reposition the wind at 0° from the front of the fly before each trial.

#### Functional imaging

All two-photon calcium imaging experiments were performed under a 20x water dipping objective (Olympus Plan Fluorite XLUMP 20x/1.0). The GCaMP7f calcium and tdTomato indicators were excited using an infrared laser (SpectraPhysics MaiTai HP) at 920nm at ∼20-40mW of power at the sample. Emitted fluorescence was detected using GaAsP photomultiplier tubes, and separated using green (GCaMP7f, 525/50) and red (tdTomato, 607/70) bandpass filters. Images were collected using ThorLabs LS 3.0 software, where 3-4 optical sections (typically 122 x 74 µm imaging window) were taken 8-12µm apart in the z plane, all within the fan-shaped body (determined using the tdTom signal). Volumes were taken at 4-8 vol/s with a 1.2µs dwell time.

#### Software control

Control of wind speed, wind position, and wind/odor state were managed by custom Python software and integrated with Fictrac ball-tracking software using an interface adapted from Python Ball Motion Tracking software from the lab of Mala Murthy. A NiDAQ board (NI PCIe-6321) and breakout board (NI BNC-2110) were used for data acquisition and communication between controllers as well as to send synchronization signals to the 2-photon microscope. Behavior data from the walking ball was collected and saved using the Fictrac image processing software at 100fps. The Fictrac timestamps (also at 100 fps) were used to trigger wind and odor changes (see experimental paradigms for timing) as well triggering 2-photon imaging (at 3s from the beginning of ball tracking).

#### Fly tethering and preparation

Flies were first anesthetized over ice in a glass scintillation vial for 5-10s, then held in a chilled aluminum “sarcophagus” (4-10°C) while being glued (McNett Aquaseal UV Fast Fix) into a custom 3D printed holder. Flies were only attached to the holder across the eyes and in front of the forelegs (episternum), so that the antennae and legs were clear and free to move. The head was tilted down to expose the back surface of the head in the saline well above. A minimal portion of this surface cuticle was removed directly above the FB to minimize tissue damage, prior to positioning the fly on the walking ball. Extracellular saline (103 mM NaCl, 3 mM KCl, 5 mM TES, 8 mM trehalose dihydrate, 10 mM glucose, 26 mM NaHCO_3_, 1 mM NaH_2_PO_4_H_2_0, 1.5 mM CaCl_2_2H_2_O, and 4 mM MgCl_2_6H_2_O, pH 7.1–7.4, osmolarity 270–274 mOsm), bubbled with carbogen (5% CO_2_, 95% O_2_) and warmed to 33°C, was continuously perfused at ∼1mL/m.

#### Experimental paradigms

Three types of experimental paradigms were used during imaging trials: 1) Closed-loop wind, open-loop odor trials (Fig. 1, Ext Data Fig. 3c,d), where fly orientation was in constant control of the wind position and odor presentation was experimentally controlled, 2) wind-shift trials (Fig. 2) where the wind position was transiently shifted during the odor period (+/- 90° at 45°/s) and 3) plume trials (Fig. 3,4), where the wind direction was in closed loop with the animal orientation and odor was in closed-loop with fly position in a fictive 2D space (see Figure 3A).

For plume experiments, the state of the odor valve was determined by an adaptive compression algorithm acting on odor concentrations based on virtual movement through a turbulent plume.

At each frame, fly position in virtual x,y space was first determined from FicTrac monitoring of ball motion. Fly coordinates in x,y, and time were used to index into a previously recorded movie of a turbulent boundary layer acetone plume at 6mm above the bed in 10 cm/s wind recorded with a UV laser light sheet^2^. Plume concentrations were normalized such that the peak concentration was 1.

Instantaneous odor concentration values were fed into an adaptive compression algorithm based on previous studies of fly behavior^3^ and of temporal encoding at the olfactory periphery^4^. Odor concentration at the *i*th frame *C_i_* was first passed through a Hill function of the form

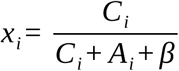

where *β* = 0.025 is the baseline threshold and *A* is an adaptation term the shifts the threshold to the right based on the recent history of odor exposures:

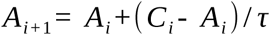

where *τ* = 500 samples (equal to 5s at 100Hz). The odor valve was turned on when *x* > 0.5 and was off otherwise.

In all experiments, we first performed 5-20 50s trials of closed-loop wind with no 2-photon imaging to acclimatize the fly to closed-loop navigation paradigm. During these trials, where wind was on for the duration of the trial, odor was presented from 20-30s, and wind speed was randomly set to one of 3 speed settings (8, 25, or 33 cm/s at the fly). After several trials, we examined behavior to see which speed was optimal for straight up-wind walking during odor and increased turning after odor offset. This speed was used for the remainder of experimental trials, although we sometimes adjusted wind speed in later trials if the fly walked less. We excluded flies that did not walk during at least 5 of these preliminary trials or only had bump activity in 2 or fewer trials.

For imaging trials, the closed-loop paradigm consisted of trial onset where wind position was randomly set to +/- 45, 135, or 0°. First the wind tube moved in closed-loop with the fly position without wind for 10s, then wind was presented at the determined speed for 15s. Odor was then presented with the wind for 15s, followed by 15s more of wind-only and 7s of no wind (total of 65s including the 3s pre-trigger time).

Experiments with plume trials also began with closed-loop speed-setting trials, followed by 10-30 plume trials that were 100s long. The fly was positioned 140 fictive mm downwind from the midline of the plume source and oriented randomly at +/- 45, 135, or 0° at the beginning of each trial. All flies were allowed to walk freely in the plume space, with no intervention if they left the plume. In this case, the fly would not receive odor until it walked back into the plume. In one fly, plume trials were run after a set of interleaved closed-loop/open-loop trials.

#### Behavioral experiments in wind tunnels

Freely walking wind tunnel experiments (Fig. 5, Ext. Data 4) were performed as described previously^2,5^. Briefly, the position and orientation of walking flies were tracked in shallow acrylic arenas using IR LEDS (850 nm, Environmental Lights) and a camera (Basler acA1920-155um). Constant laminar airflow was maintained at ∼12 cm/s. Odor pulses of 1% apple cider vinegar (Pastorelli) were delivered at 1 Hz (500 ms duration) using solenoid valves (LHDA1233115HA, Lee Company) mounted directly below the wind tunnel. For GtACR1 inactivation experiments, blue light was delivered using a panel of LEDs (470nm, Environmental Lights) at an intensity of 61 µW/mm^2^ (measured at 460 nm). For high light conditions, the intensity was set to 76 µW/mm^2^ (measured at 460 nm). Flies were briefly anesthetized with ice for 30-60 seconds to load them into the wind tunnels.

## Data Analysis

### All analysis was performed in MATLAB (Mathworks, Natick, MA)

#### Processing of walking ball data

For walking ball data, angular velocity and forward velocity were calculated as the change in rotation of the ball around the z and x axes, respectively. For closed-loop wind trials, heading was reported as the position of the wind tube, which was controlled by the change in z rotation, but would also account for any error in the control. For plotting and bump detection, heading, angular velocity, and forward velocity signals were smoothed using a sliding averaging window of 100 frames (1s). For linear filter calculations (Fig. 3e, 4b,c,), we used unfiltered values as described above.

Speed was calculated as the velocity in direction of heading based on the fictive fly path derived from integrating all three rotational velocities to map position on to a 2D plane^1^. X and Y position of the fly were calculated as the cumulative sum of the speed multiplied by the sine or cosine, respectively, of heading. For plume trials, X and Y position values were calculated online to enable closed-loop odor control. Upwind velocity (Fig. 4, Ext. Data Fig. 1a,3b) was calculated as the cosine of heading multiplied by speed.

Straightness (Ext. Data Fig. 1a,3e) was computed as the ratio of displacement (the distance of a straight line from the beginning and end points of a window) to path length (the sum of all displacements within the window) over a sliding 1.5s window. This value was then squared. Persistent walking duration (Ext. Data Fig. 1b,c) was determined as the amount of time walking in the goal direction before a heading deflection of more than ±45° in trials with bump activity (see below bump detection methods) and pooled across flies. Goal direction (Ext. Data Fig. 1b) was determined as the heading at odor OFF for odor walks, and heading at trial start for baseline. The distribution of durations was compared to the distribution of straight walking from the trial onset (Ext. Data. Fig. 1c), using identical criteria as post odor runs.

Exponential curves were fit to these distributions using a non-linear least-squares process. The curve was defined as:

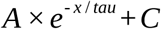

where A is the amplitude of the starting value, tau is the decay rate, and C is the offset. Optimal fit was determined by minimizing mean squared error. A 95% confidence interval (CI) for tau was given by:

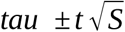

where *t* is computed using the inverse of Student’s *t* cumulative distribution function, and *S* is a vector of the diagonal elements from the estimated covariance matrix of the coefficient estimates, (*X^T^X*)^−1^*s*^2^, where *X* is the design matrix, *X^T^* is the transpose of *X*, and *s*^2^ is the mean squared error (using the Curve Fitting Toolbox from Mathworks).

#### Processing of imaging data

For fluorescence extraction, all image volumes were first reduced to single planes as maximum intensity projections across each z plane. Maximum intensity projections were highly correlated with mean projections, but had slightly higher signal-to-noise. We then concatenated projections for all trials of an experiment, and used the NoRMCorre algorithm^6^ to correct for rigid motion across both red and green channels. We drew ROIs by hand in ImageJ, using a maximum intensity projection across all frames of a single trial (typically the first trial) on the red tdTOM channel. We then imported these ROIs into Matlab, and aligned them to subsequent images using register_rois2^7^. For both VT062617 and 52G12 imaging, we drew ROIs for each of 8 putative columns in the dorsal layer of the FB. A baseline value for each ROI was calculated as the mean of the lowest 3% of fluorescent values across all trials. ΔF/F was then calculated by subtracting this baseline value from the green fluorescence values and dividing them by the baseline value. Fluorescence timeseries data was synchronized to the timestamps of behavior data using the 3s trigger time.

#### Analysis of neural encoding properties

To detect activity bumps, we used a threshold-based algorithm, where the threshold for ΔF/F was set to 0.5 standard deviations above the mean ΔF/F of the entire experiment. A minimum bump duration was set at 3s and bumps with gaps <3s from a consecutive bump were merged. Using this detector, we classified samples of behavior data as either bump or no-bump samples, and detected bump onset and offset times. Only flies with more than 2 trials showing bump activity were included.

To visualize the relationship of bump activity to odor (Fig. 1e), the probability of bump activity of all trials was shown aligned with odor onset for all closed-loop trials with at least one bump. Bump events were identified using the previously defined threshold and converted to a binary time series indicating bump presence. For visualization, trials were pooled across flies to generate heat maps, and mean bump probability was computed across trials. To generate distributions of post-odor bump durations (Fig. 1f,g, and Ext Data. Fig 1i), we analyzed all bumps that began during the odor period and ended after odor OFF.

To analyze how behavior changed during neural bump events (Fig 1h, Ext. Data Fig. 1j,k), bumps were found in all closed-loop trials and behavioral metrics were computed over the full bump duration and over a paired pre-bump window of the same duration immediately preceding onset. Behavioral metrics were calculated within each window (standard deviation of unwrapped heading or mean angular speed). Bump events were also categorized by onset time relative to odor presentation (pre-odor, during odor, or post-odor), and all bump–pre pairs for which a pre-bump window was available in the data were pooled across flies within each category. Paired two-tailed t-tests were used to compare bump versus pre-bump behavioral values within each category.

To analyze behavior associated with bump offset or bump onset (Fig. 1i,j, Fig. 3d, Ext. Data Fig. 2d), we selected bumps that turned ON during odor presentation. For the bump offset analysis (Fig. 1i,j, Fig. 3d), we only included bumps that also turned OFF after odor OFF. We then computed locomotor parameters (angular speed and forward velocity) associated with these periods as described above. For bump offset analysis, we also computed goal deviation as the absolute value of the difference between the absolute value of heading at each time point after odor OFF and a goal heading (θ_goal_), defined as the mean heading during the odor period. We then aligned each behavioral parameter by the time of bump OFF or ON and took the mean across all trials from all flies. For comparison (control), we aligned behavioral data to the midpoint of each bump. Statistical comparisons were made over a window from 0-2.5s after bump off (vs bump midpoint) for angular speed, and for a window from 0-5s after bump off (vs bump midpoint) for goal deviation. For bump onset (Ext. Data Fig. 2d), we compared mean forward velocity over a window -7 to -4 s before bump onset to a window -1.5 to 1.5 s from onset of each bump. A paired two-sample t-test was used for all comparisons.

To compute bump width in Ext Data Fig. 1g, we aligned each bump sample to the ROI with the peak ΔF/F, and then wrapped the other ROI values to the 8 column space using the modulus function in Matlab. Aligned bumps were averaged across trials and flies.

To compute bump position in Fig. 2, we first smoothed fluorescence traces using a 1s sliding window in the time domain and a two-column window in the spatial domain. We then computed bump position as the maximum ROI (or FB column) for each sample where a bump was detected using our bump finder algorithm above. Positions were then unwrapped to show absolute column shifts. We computed correlations between bump position and heading for all bump instances (Fig. 2b, top) and averaged across each fly with bump activity in at least 5 trials (Fig. 2b, bottom) using Pearson’s correlation coefficient.

To compute bump position shifts associated with a rotation in wind direction, in Fig. 2c,d, we analyzed only trials in which a bump was present before and during the shift. Wind driven heading shifts were detected when mean angular velocity during or up to 2s after the shift was greater than a time window of equal duration before the shift. Heading and position for right shifts were multiplied by -1 so all shifts could be directly compared on the same axes. Heading and bump position were both converted to degrees to facilitate comparison. Displacements were calculated over 2s time windows before and after the wind shift onset. For these trials, heading was extracted directly from integrated ball yaw rotations, rather than the wind stimulus position, as in trials without a wind shift.

To compute linear filters relating odor experience and locomotor parameters to neural activity in Fig. 3e, Fig. 4b,c, Ext. Data Fig. 2a-c, we first computed bump amplitude, forward velocity, and angular velocity as described above. The first 400 samples of each trace were discarded as the bump amplitude typically contained an artifact due to shutter opening. The remaining traces were concatenated to form a single vector for each parameter and all traces were mean subtracted. Forward velocity and absolute angular velocity were normalized to have unit standard deviation. We then computed the cross-correlogram between bump amplitude and odor, forward velocity, and absolute angular velocity in the frequency domain.

To account for correlations between odor and locomotor parameters, we adapted Methods from^9^ to account for correlations between odor, forward velocity, and angular velocity. Briefly, we computed the complex autocorrelation matrix A at each of 256 frequencies from 0.2 to 5 Hz of the odor o(t), forward velocity f(t), and absolute angular velocity a(t) traces:

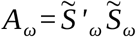

where

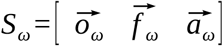

We then computed the pseudo-inverse of *A* for each frequency *ω* by taking the singular value decomposition:

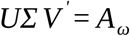

and inverting the diagonal terms of *Σ* with a regularization parameter *λ* that was set to 0.0001.

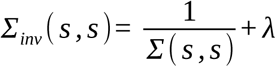

The decorrelated filter was equal to:

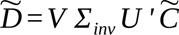

Where 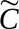 is the Fourier transform of the raw cross-correlogram for each parameter:

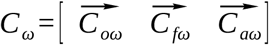

To compare filters from different fly lines in Fig. 4b,c, we normalized all filters by dividing by the standard deviation of the filter and subtracting minimum standard deviation values. To compare half-widths of filters, we first smoothed each filter using a sliding averaging window of 1s. We then found the half-prominence width of peaks determined as local maxima where the prominence was a minimum of 1.5 times the mean for each fly (Matlab findpeak function, with defined prominence properties). To compare angular velocity filters, we found the area under the curve for each fly using a trapezoidal estimation for each data point (Matlab trapz function).

To compute the correlation between upwind velocity and bump activity on a trial-by-trial basis in Fig. 4d,e, we measured upwind velocity during the odor period for each trial. We then computed Pearson’s correlation coefficient between these metrics and the mean ΔF/F for the same periods and trials. To assess significance, we compare these to correlation coefficients between metrics and ΔF/F for shuffled trials (null condition).

#### Analysis of freely walking wind tunnel behavior data

Freely walking behavioral data were preprocessed as described previously^2,5^:

Upwind walking probability was determined by binarizing the orientation data depending on whether the orientations were inside or outside of a 90° cone centered on upwind. This binarized data was further processed in three ways to evaluate persistence and evidence accumulation. First, the binarized data were averaged at each timepoint to compute the upwind walking probability over time (Fig. 5b,d,e). Second, the upwind walking probabilities over time were averaged between 2 s and 6 s following odor onset to compute peak responses (Fig.5c). Third, the binarized data between 0 and 10 s following odor offset were averaged across trials for each fly to compute post-odor upwind orientation probabilities (Fig. 5f,g).

#### Modeling persistent goal-directed navigation in an odor plume

We implemented a three-state behavioral model to simulate fly navigation in a natural odor plume environment. At each time step, the agent occupied one of three discrete states: baseline walking, goal-directed upwind walking, or local search. Transitions between states were governed by odor inputs and probabilistic rules inspired by our observed neural dynamics. The agent received odor concentration values sampled from the same measured plume used in imaging experiments described above, and the raw odor signal was passed through the same adaptive filter and compression function with *β* = 0.025 and *τ _A_* = 5 *s*.

When the compressed odor signal C*_t_* exceeded a threshold (θ=0.5), the agent transitioned from baseline or search into the goal-directed state. The agent transitioned from the goal state to the search state with a probability defined by the persistence time constant τ:

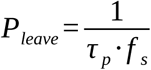

where *f*_s_ is the simulation frame rate (15 Hz). From the search state, it could return to the goal state if odor reappeared, or stochastically transition back to baseline with a fixed time constant (6.7s).

Behavior in each state was described by a modified version of the model in^26^. Briefly, locomotion was controlled by a network of 5 descending neuron-like units that receive independent Gaussian noise 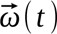 and interact through a connectivity matrix M. *U* _3_ drives stopping while the remaining units drive forward and angular velocity through their sums and differences:

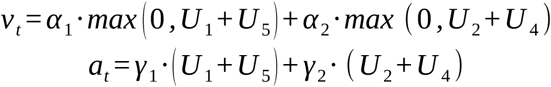

We adjusted the parameters α_1_,α_2_, γ_1_, γ_2_ to approximately match the distribution of forward and angular velocities measured on our walking ball during boundary layer plume navigation (Fig. 6c).

|  | Walking | Flight |
| --- | --- | --- |
| $\alpha_1$ | 0.7°/s | 40°/s |
| $\alpha_2$ | 0.3°/s | 30°/s |
| $\gamma_1$ | 0.1mm/s | 0.9mm/s |
| $\gamma_2$ | 0.2mm/s | 0.9mm/s |

To assess and optimize these parameters, we compared measures of their respective behavior distributions (Fig. 6c). Forward and angular velocity samples were extracted from raw behavioral time series in real flies and model simulations. For real data, closed-loop trials were analyzed over the pre-odor segment, selecting for trials averaging > 1 mm/s. Forward velocity was restricted to v > 0 mm/s, and angular velocity was analyzed as magnitude (|ω|). Model outputs were processed using the same criteria.

Distributions were visualized using normalized histograms, while statistical analyses were performed on unit-level means (flies for real data, trials for model data). Mean differences were assessed using Welch’s 95% confidence interval.

#### The matrix *²* has the structure

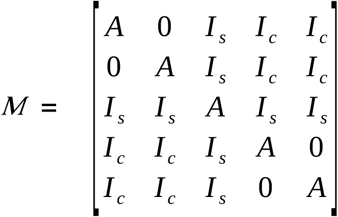

During the baseline and goal states, parameters were *A* = 0.8, *I _s_* = - 0.03, *I _c_* = 0. During the search state, *I _c_* = - 0.035. The inhibition between the two sides present during the search state reduces forward velocity and increases both angular velocity and the duration of runs of turns in the same direction, as described in^26^. During the goal state, a sinusoidal function of wind direction was added to the locomotor units to promote an upwind heading and the stop unit was inhibited to promote greater locomotion as in^26^:

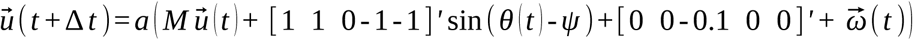

Activity in each unit was bounded by a sigmoidal activation function that prevents runaway excitation:

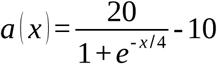

To assess how the persistence parameter *τ _p_* influences navigation success, we ran the model across a range of *τ _p_* values in the near-surface (boundary layer) air plume described above^3^ (Fig. 6d,e). The plume environment measured 300.44mm x 159.84 mm. For each *τ _p_*, 500 trials were simulated with randomized start positions downwind from the source by 222±12mm and centered along the midline of the plume’s long axis, ±100mm. Success was quantified by calculating the proportion of time the agent’s position fell within a 15 mm radius of the odor source.

To visualize where the agent engaged in each behavioral state (Fig. 6d), we computed spatial heatmaps showing the probability of occupying the baseline, goal-directed, or search state across the plume environment. The agent’s position and corresponding state label were divided by 15mm x 15mm bins, and the relative frequency of each state within each bin was calculated. These probability maps were smoothed using a Gaussian filter (σ = 15mm) to highlight spatial patterns.

### Immunohistochemistry

To perform immunohistochemistry, dissected brains were fixed in 4% paraformaldehyde (dissolved in PBS) for 15 minutes, washed 3x in PBS, incubated in 5% normal goat serum (dissolved in PBST) for 60 minutes, incubated overnight in primary antibody solution (dissolved in 5% normal goat serum), washed 3x in PBST, incubated overnight in secondary antibody solution (dissolved in 5% normal goat serum), washed 3x in PBST, and washed 3x in PBS. The primary antibody solution contained chicken anti-GFP (Fisher Scientific RRID:AB_1074893) 1:50, mouse anti-nc82 (DSHB RRID:AB_2314866) 1:50, and rabbit anti-dsRed (Clontech 632496) 1:500. The secondary antibody solution contained Alexa488-conjugated goat antichicken (Fisher Scientific RRID:AB_2534096) 1:250, Alexa633-conjugated goat anti-mouse (Fisher Scientific RRID:AB_2535719) 1:250, and Alexa568-conjugated goat anti-rabbit (Fisher Scientific RRID:AB_2576217) 1:250. To image the brains, the brains were mounted on microscope slides, immersed in vectashield (Vector Labs H- 1000), sealed with coverslips, and then imaged using a 20× objective (Zeiss W Plan-Apochromat 20×/1.0 DIC CG 0.17 M27 75 mm) on a Zeiss LSM 800 confocal microscope at 1.25 μM depth resolution.

**Table 1:** Reagents and Resources.

| Resource | Source | Identifier |
| --- | --- | --- |
| **FLY LINES** |  |  |
| VT062617-Gal4 | VDRC | VDRC: 206875 |
| 52G12-Gal4 | BDSC | RRID: BDSC_49581 |
| Empty-Gal4 | BDSC | RRID: BDSC_68384 |
| w<sup>1118</sup> 5905 | Alvarez-Salvado et al. 2018 |  |
| UAS-GCaMP7f,UAS-tdTomato | BDSC/Nagel lab | RRID: BDSC_80906, 36327 |
| w<sup>1118</sup>,s/CyO;UAS-GtACR1-EYFP/TM6b | Claude Desplan |  |
| VT062617AD/[CyO,tb];VT037223DB/TM6b | Tanya Wolff | Janelia: SS63158 |
| p65ADZp, ZpGAL4DBD (Empty Split-Gal4) | BDSC | RRID: BDSC_79603 |
| **SOFTWARE** |  |  |
| Python |  | Version 3.8 |
| MATLAB | Mathworks | Version R2023a |
| Image J |  | Version 1.53t |
| FicTrac |  | Version 2.0 (build date: Nov 2 2018) |
| Pybmt | Mala Murthy lab |  |
| CalmAn |  |  |
| Register_roi2 | Jason Moore |  |

## Data Availability

Data will be made publicly available upon publication at Zenodo.

## Code Availability

Model code is available at https://github.com/nagellab/Kathmanetal2025.

## Acknowledgements

The authors would like to thank Mala Murthy and her lab for providing modified FicTrac code (Pybmt) and advice on closed-loop behavior, John Crimaldi and his lab for plume data, and Margot Elmaleh in Michael Long’s lab for advice on 3D printing. Claude Desplan’s lab provided GtACR1 flies. David Schoppik provided assistance with 2-photon microscopy. Christina May generated imaging stocks and provided support for 2-photon microscopy. Emily Hao provided assistance with behavioral experiments. Jason Moore provided assistance with imaging analysis. Jonathan Victor provided advice on regularizing the filter decorrelation. Michael Long, Jonathan Victor, Bard Ermentrout, Floris van Breugel, and members of the Nagel and Schoppik labs provided input on the manuscript. This work was supported by NINDS Brain Initiative NS127129, NIDCD DC017979, and NSF 2014217 Odor2Action to K.I.N.

## Author Contributions

N.D.K. and K.I.N designed the study. N.D.K. developed the closed-loop wind and odor apparatus, developed all all hardware and software related to the apparatus, and collected and analyzed all behavior and imaging data using this apparatus. A.J.L designed, ran, and analyzed freely walking behavioral data and contributed analysis methods and code. K.I.N. wrote code for the filter analysis. J.D.F. obtained confocal data. N.D.K. and K.I.N. wrote the paper with input from A.J.L.

## Competing Interest Declaration

The authors declare no competing interests.

## Extended Data

**Ext. Data Fig. 1:**
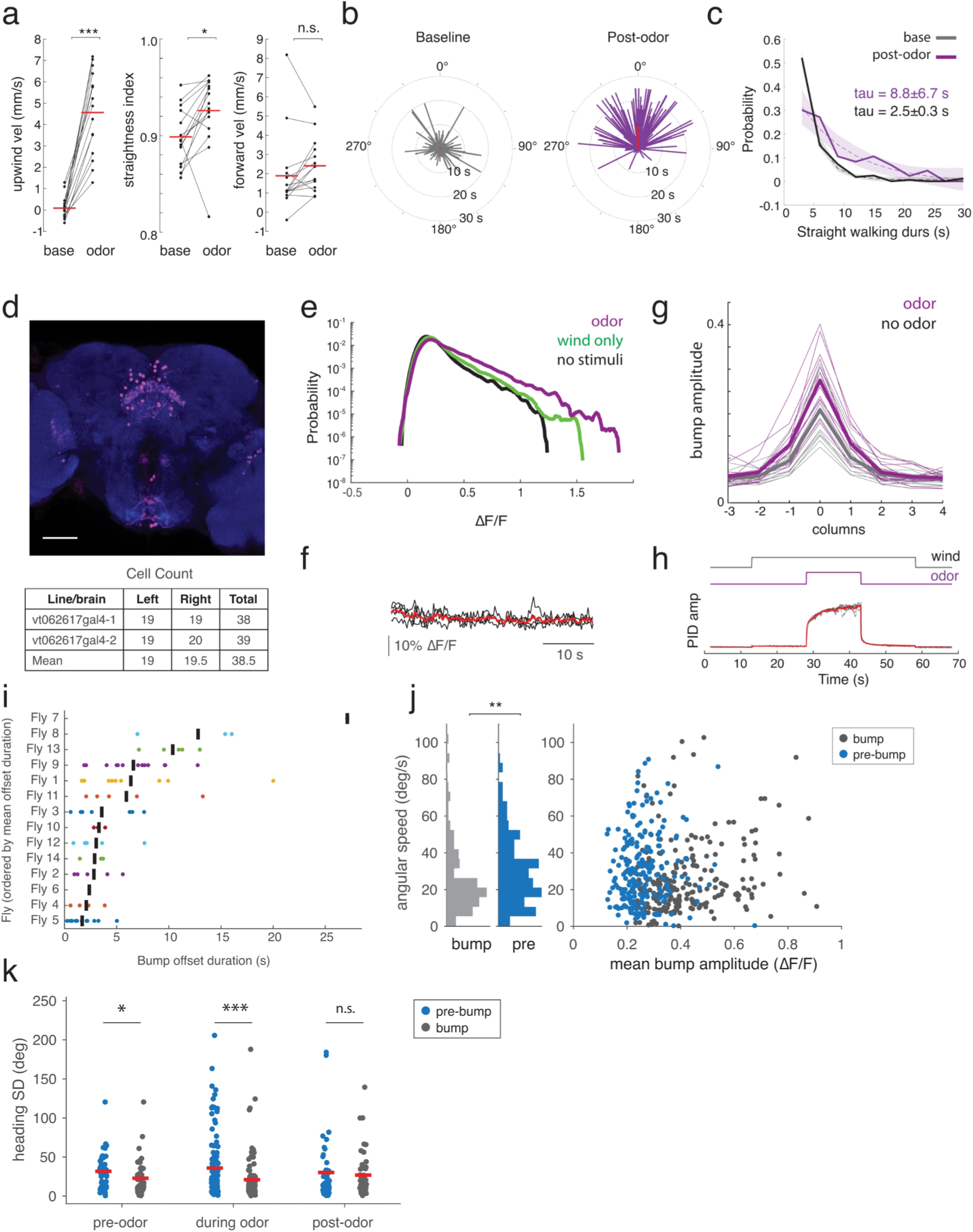
Additional data on VT062617 FB local neuron responses. **a**: Characteristics of virtual trajectories during baseline versus odor periods for all flies (N=15). Each line represents the mean for one fly. Mean upwind velocity and straightness are significantly higher during odor than baseline (paired t-test, p =0.001, p=0.035). Forward velocity is not significantly different (p=0.43). **b.** Heading distribution of persistent walking during baseline (left) and odor OFF (right) periods for all trials with bump activity (n = 135 trials from N=14 flies). Mean resultant vector during post-odor (red) is biased upwind (mean: -1.5°). The length of each line represents the duration the animal walked straight after the respective time (odor: odor OFF, baseline: trial start). c. Distribution of persistent straight walking durations after odor (purple) and during baseline (grey) for each fly (N=14). Exponential curves (dashed) with functional confidence intervals (shaded) were fit to the data, providing exponential decay values (tau_odor_ = 8.8±6.7s, tau_base_ = 2.5±0.3s). **d**: Top: Confocal image of VT062617>GCaMP brain, stained for GCaMP (pink) and nc82 (blue). Fluorescence signals were imaged from the dorsal (output) projections of FB local neurons. Scale bar is 50µm. Bottom: table of cell body counts for left and right hemispheres in two brains. **e**: Distribution of ΔF/F values across all recordings in the presence of odor (purple), during wind only (green) and in the absence of stimuli (black). Total of 328 trials from 15 flies for each condition. **f**: Average bump amplitude in trials with no wind or odor stimuli, but motor still in closed-loop with heading (n=3-7 trials, N=3 flies). **g**: Mean bump profile across columns. Thin lines represent mean across trials (9 to 52 trials for each fly) and thick lines represent mean across flies (N=14 flies with both odor and no-odor imaging data). **h**: Photoionization detector (PID) recording of odor concentration during typical wind and odor pulse trials (n=6 trials, black: individual trials, red: mean across trials) at the location of the fly. Wind and odor control signals above. **i**: Mean bump persistence distributions of individual flies are significantly different from one another (one-way ANOVA, p = 2.29e-10, n = 82, N = 14). Each row shows all bump persistence times for a single fly, with the mean indicated by a black line. Flies are sorted along the y axis by mean duration. **j:** Mean angular speed (y-axis) versus bump amplitude (x-axis) for all bumps (222 bumps from 14 flies, blue) and for equivalent durations before each bump. Marginal distributions at left. Angular speed during bumps is significantly lower than during pre-bump periods (p=0.01). **k**: Standard deviation of unwrapped heading was smaller during bump versus pre-bump periods for bumps starting during the pre-odor (n=52 bumps, p=0.03), and odor (n=123 bumps, p=3.6e-6) phases, but not for bumps starting during the post-odor period (n=47 bumps, p=0.60). The smaller reduction observed for bumps starting in the post-odor period may reflect the fact that pre-bump periods occur during odor in this case.

**Ext. Data Fig. 2:**
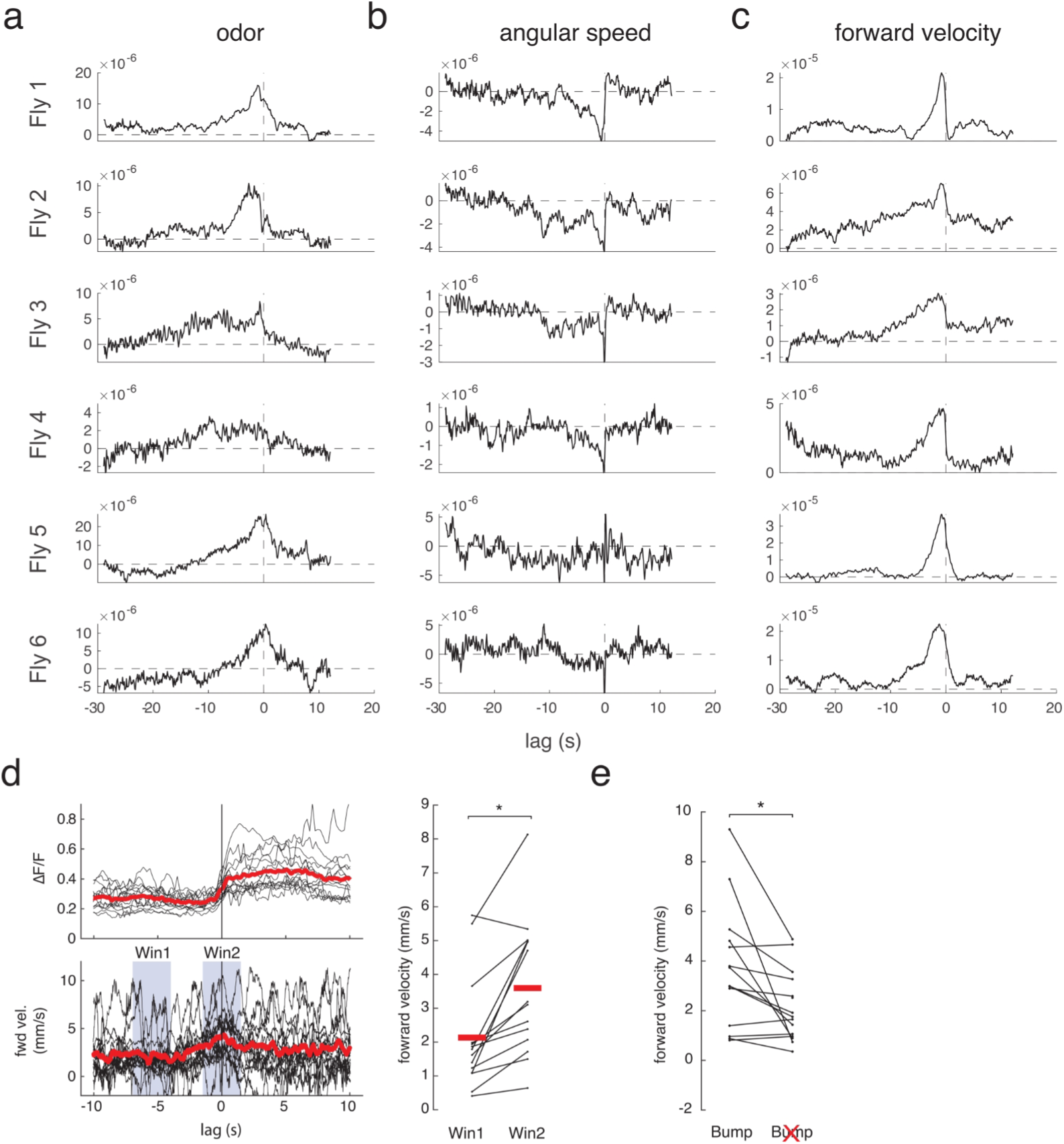
Additional data on plume related activity and bump onset. **a**: Linear filters for odor (**a**), angular speed (**b**), and forward velocity (**c**) for each fly. **d**: Analysis of locomotor activity relative to bump onset in single odor pulse trials. Left: lower trace shows forward velocity aligned by bump onset (upper), which peaks at bump onset. Red traces represent means across flies (N=14 flies). Right: Mean forward velocity increases during bump onset (pre-onset mean=2.13, post-onset mean=3.60, paired t-test p=0.046, blue boxes indicate analyzed periods.) **e**: Mean forward velocity during odor is higher in samples with a bump than without (paired t-test, p=0.032, N=14).

**Ext. Data Fig. 3:**
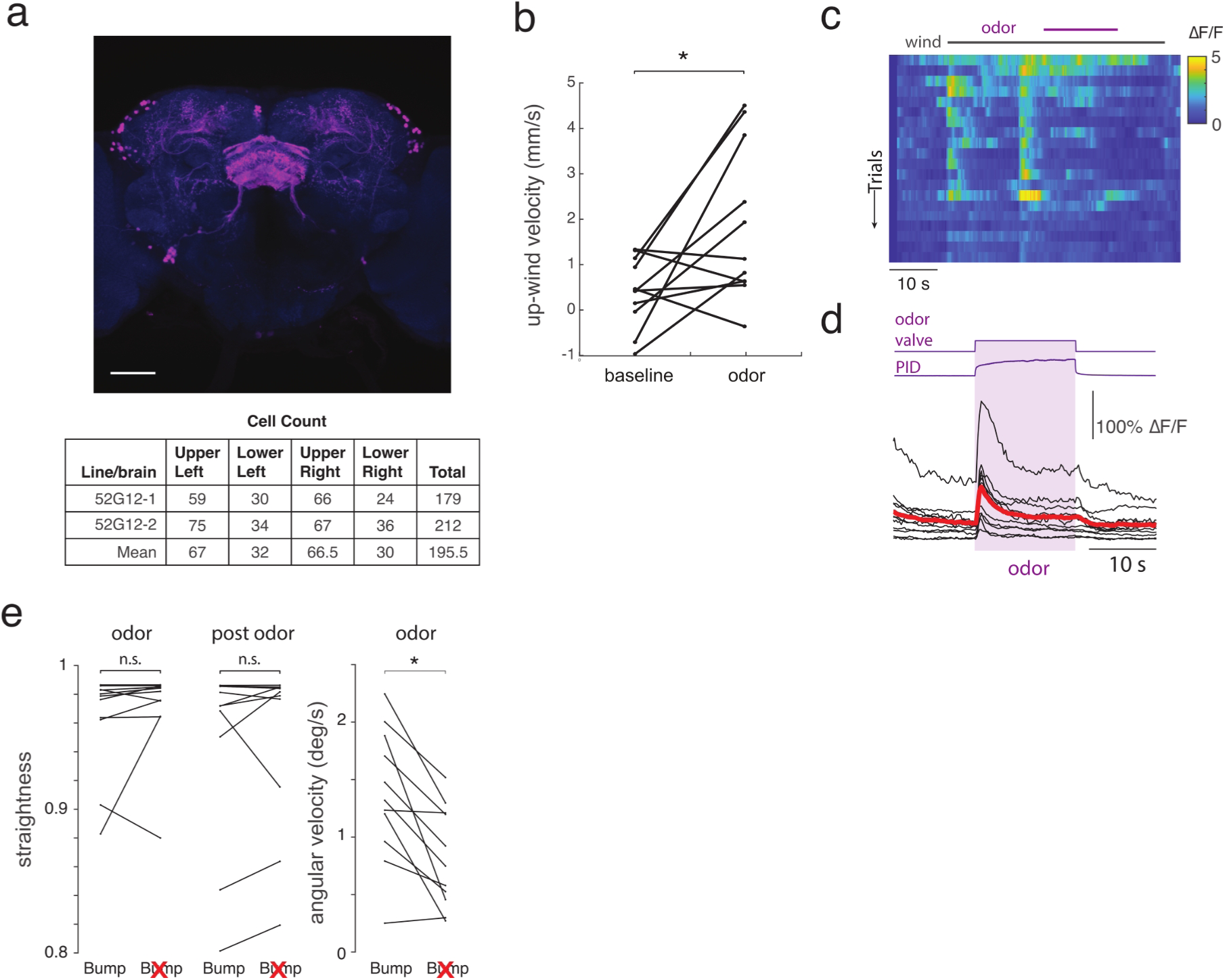
Additional data on responses of ventral local neurons labeled by 52G12. **a**: Confocal image of 52G12>GCaMP brains, stained for GCaMP (pink) and nc82 (blue). Scale bar is 50µm. Bottom: table of cell body counts for left/right/upper/lower quadrants of two brains imaged. **b**: Mean upwind velocities of expressing GCaMP in 52G12 neurons during odor versus baseline (N=11 flies, p=0.018). **c**: Mean activity in 52G12 neurons over time for all trials for one fly. Note transient responses to wind and odor onset. **d**: Time course of activity relative to measured odor time course (purple box) for all recorded flies. Black: mean for each fly (n=6 to 30 trials each), red: mean across flies (N=11 flies). **e**. Left: Mean straightness for 52G12 flies (N=11) in the odor and post-odor periods is not significantly different in the presence versus absence of a bump (paired t-test, during odor p=0.966, post-odor p=0.893). Right: Angular speed during odor is significantly higher in the presence of a bump (paired t-test, p=0.029)

**Ext. Data Fig. 4:**
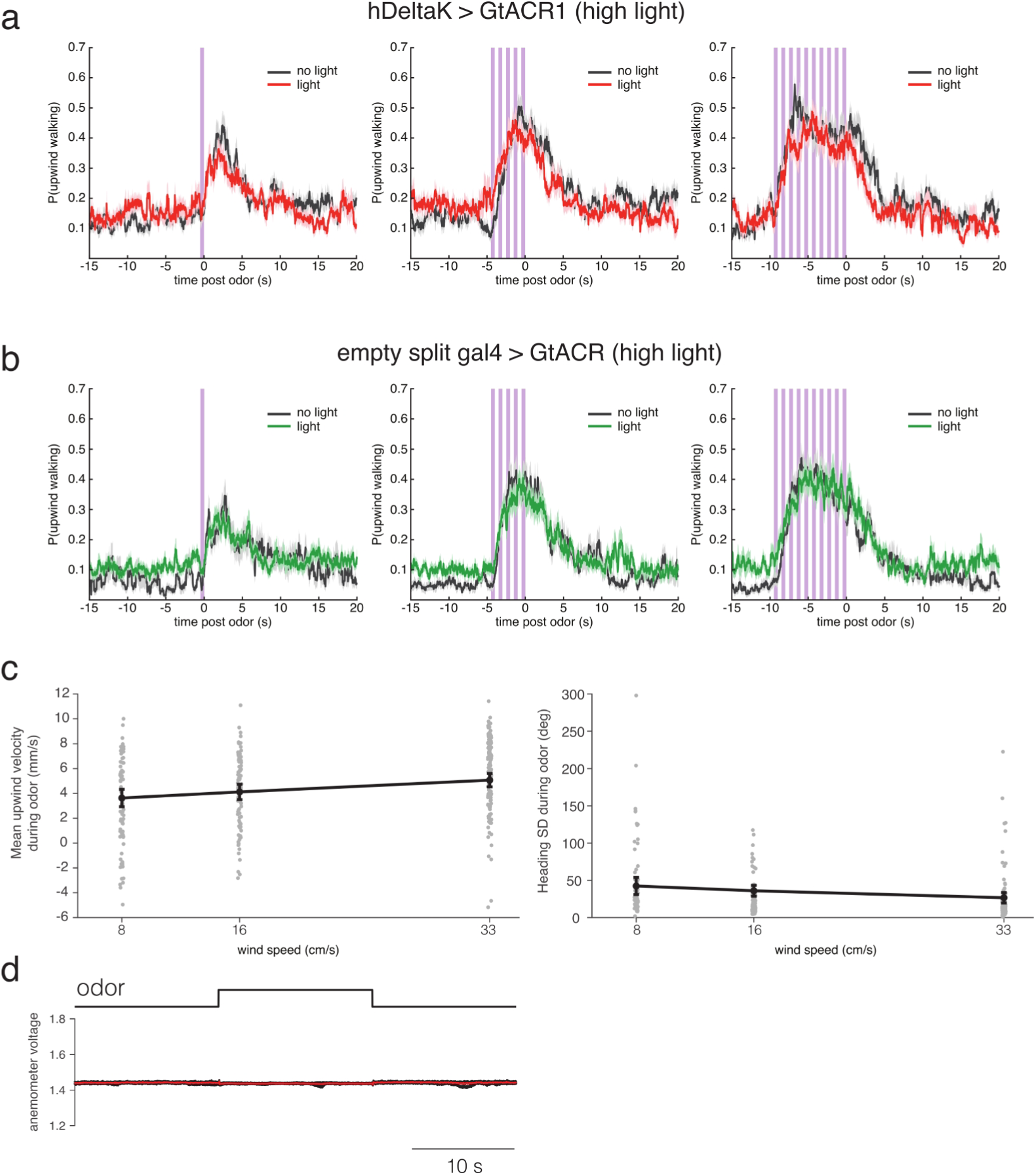
Additional data on silencing of hΔK neurons. **a**: Probability of upwind walking in hΔK split > GtACR flies responding to 1, 5, or 10 pulses of odor (500 ms at 1Hz) with light OFF (black, 1 pulse: n=287 trials from N=23 flies; 5 pulses: n=271 trials from N=24 flies; 10 pulses: n=297 trials from N=24 flies) or light ON (red, 1 pulse: n=362 trials from N=23 flies; 5 pulses: n=367 trials from N=24 flies; 10 pulses: n=322 trials from N=24 flies). Light level for these experiments was 76 µW/mm^2^. We observed a decrease in upwind walking probability after odor OFF on light ON trials for the 10 pulse condition. **b**: Probability of upwind walking in empty-split > GtACR flies responding to the same stimuli with light OFF (black, 1 pulse: n=150 trials from N=26 flies; 5 pulses: n=234 trials from N=28 flies; 10 pulses: n=213 trials from N=29 flies) and light ON (green, 1 pulse: n=357 trials from N=26 flies; 5 pulses: n=324 trials from N=28 flies; 10 pulses: n=358 trials from N=29 flies). **c**: Dependence of upwind tracking on wind speed during walking ball trials. Left: upwind velocity during odor showed a positive correlation with wind speed (rho =0.29, p = 3.2e-07) but no significant relationship to wind speed by ANOVA (p=0.283). Right: standard deviation of unwrapped heading during odor showed a negative correlation with wind speed (rho =-0.32, p = 2.4e-08) but no significant relationship to wind speed by ANOVA (p=0.427). Gray dots represent means of single trials; black circles represent mean across trials. **d:** anemometer trace (6 trials in black, mean in red) measured at the location of the fly during an odor pulse.

**Extended Data Table 1:** Statistics for Figure 5. Significant p values are bolded.

| Figure | Comparison | Sample size | Statistical test | p value |
| --- | --- | --- | --- | --- |
| Fig. 5b-c | P(upwind orientation) for 1, 4, 7, or 10 odor pulses | N<sub>1 pulse</sub> = 30 flies, 246 trials<br>N<sub>4 pulses</sub> = 28 flies, 226 trials<br>N<sub>7 pulses</sub> = 29 flies, 213 trials<br>N<sub>10 pulses</sub> = 30 flies, 246 trials | N/A | N/A |
| Fig. 5d | VT062617 > GtACR P(upwind orientation) for 1, 5, or 10 odor pulses with or without light | Light OFF:<br>N<sub>1 pulse</sub> = 30 flies, 381 trials<br>N<sub>5 pulses</sub> = 30 flies, 418 trials<br>N<sub>10 pulses</sub> = 30 flies, 463 trials<br>Light ON:<br>N<sub>1 pulse</sub> = 30 flies, 405 trials<br>N<sub>5 pulses</sub> = 30 flies, 422 trials<br>N<sub>10 pulses</sub> = 30 flies, 450 trials | N/A | N/A |
| Fig. 5e | Empty-Gal4 > GtACR P(upwind orientation) for 1, 5, or 10 odor pulses with or without light | Light OFF:<br>N<sub>1 pulse</sub> = 26 flies, 261 trials<br>N<sub>5 pulses</sub> = 26 flies, 246 trials<br>N<sub>10 pulses</sub> = 28 flies, 318 trials<br>Light ON:<br>N<sub>1 pulse</sub> = 26 flies, 294 trials<br>N<sub>5 pulses</sub> = 28 flies, 310 trials<br>N<sub>10 pulses</sub> = 27 flies, 292 trials | N/A | N/A |
| Fig. 5f | Mean P(upwind orientation) for 1, 5, or 10 odor pulses with or without light, for VT062617 > GtACR and empty-Gal4 > GtACR1 flies | VT062617 > GtACR:<br>N<sub>1 pulse</sub> = 26 flies<br>N<sub>5 pulses</sub> = 26 flies<br>N<sub>10 pulses</sub> = 26 flies<br>empty-Gal4 > GtACR:<br>N<sub>1 pulse</sub> = 25 flies<br>N<sub>5 pulses</sub> = 27 flies<br>N<sub>10 pulses</sub> = 28 flies | Bonferroni-corrected ranksum test ($\alpha = 0.017$) | **VT062617 > GtACR 1 pulse:**<br>**p < 0.0001**<br>**VT062617 > GtACR 5 pulses:**<br>**p = 0.0046**<br>**VT062617 > GtACR 10 pulses:**<br>**p = 0.0004**<br>empty-Gal4 > GtACR 1 pulse:<br>p = 0.474<br>empty-Gal4 > GtACR 5 pulses:<br>p = 0.046 |
|  |  |  |  | empty-Gal4 ><br>GtACR 10 pulses:<br>p = 0.399 |
| Fig. 5g | Mean P(upwind orientation) for 1, 5, or 10 odor pulses with or without light, for hΔK split > GtACR and empty split-Gal4 > GtACR1 flies | hΔK split > GtACR:<br>N<sub>1 pulse</sub> = 23 flies<br>N<sub>5 pulses</sub> = 24 flies<br>N<sub>10 pulses</sub> = 24 flies<br>empty split-Gal4 > GtACR:<br>N<sub>1 pulse</sub> = 26 flies<br>N<sub>5 pulses</sub> = 28 flies<br>N<sub>10 pulses</sub> = 29 flies | Bonferroni-corrected ranksum test (α = 0.017) | VT062617 > GtACR 1 pulse:<br>p = 0.276<br>VT062617 > GtACR 5 pulses:<br>p = 0.065<br>**VT062617 > GtACR 10 pulses:**<br>**p = 0.0079**<br>empty-Gal4 > GtACR 1 pulse:<br>p = 0.764<br>empty-Gal4 > GtACR 5 pulses:<br>p = 0.991<br>empty-Gal4 > GtACR 10 pulses:<br>p = 0.417 |

## Notes

### Competing Interest Statement

The authors have declared no competing interest.

### Summary of Updates

We have updated analyses in Figures 1 and 3 and moved data on the neural and behavioral responses to wind shifts to a new main Figure 2.

